# The development of hepatic steatosis depends on the presence of liver-innervating parasympathetic cholinergic neurons in mice fed a high-fat diet

**DOI:** 10.1101/2023.11.03.565494

**Authors:** Jiyeon Hwang, Junichi Okada, Li Liu, Jeffrey E. Pessin, Gary J. Schwartz, Young-Hwan Jo

## Abstract

Hepatic lipid metabolism is regulated by the autonomic nervous system of the liver, with the sympathetic innervation being extensively studied, while the parasympathetic efferent innervation is less understood despite its potential importance. In this study, we investigate the consequences of disrupted brain-liver communication on hepatic lipid metabolism in mice exposed to obesogenic conditions. We found that a subset of hepatocytes and cholangiocytes are innervated by parasympathetic nerve terminals originating from the dorsal motor nucleus of the vagus. The elimination of the brain-liver axis by deleting parasympathetic cholinergic neurons innervating the liver prevents hepatic steatosis and promotes browning of inguinal white adipose tissue (ingWAT). The loss of liver-innervating cholinergic neurons increases hepatic *Cyp7b1* expression and fasting serum bile acid levels. Furthermore, knockdown of the G protein-coupled bile acid receptor 1 gene in ingWAT reverses the beneficial effects of the loss of liver-innervating cholinergic neurons, leading to the reappearance of hepatic steatosis. Deleting liver-innervating cholinergic neurons has a small but significant effect on body weight, which is accompanied by an increase in energy expenditure. Taken together, these data suggest that targeting the parasympathetic cholinergic innervation of the liver is a potential therapeutic approach for enhancing hepatic lipid metabolism in obesity and diabetes.

## Introduction

Metabolic dysfunction-associated fatty liver disease (MAFLD), particularly its progressive form known as metabolic dysfunction-associated steatohepatitis (MASH), is now the leading cause of liver transplantation in Western countries [1]. MAFLD is characterized by the accumulation of excessive amounts of fat in the liver, which is determined by multiple factors including the uptake of fatty acids by hepatocytes, *de novo* lipogenesis (DNL), mitochondrial and peroxisomal fatty acid oxidation, and lipid export from the liver [2, 3]. Results from recent studies support important roles for the autonomic nervous system in mediating multiple critical hepatic lipid metabolism pathways . For example, MAFLD both in mice and humans [4] is associated with significant plasticity in the sympathetic innervation to the liver, including swollen axonal varicosities, nerve fiber retraction, and abnormal axonal sprouting. The vagus nerve also exerts a significant parasympathetic influence on the regulation of hepatic lipid metabolism. For example, activation of central leptin receptors reduces hepatic fat accumulation, which requires surgically intact hepatic vagal innervation in lean and high-fat fed rodents [5]. Similarly, the anti-steatotic effect of leptin is dependent on vagal innervation in humans [6]. In addition, central infusion of oleic acid suppresses triglyceride-rich very low-density lipoproteins (VLDL-TG) secretion through hepatic vagal innervation in rats [7]. The hepatic vagus nerve regulates the development of MAFLD via altering hepatic inflammation [8]. However, a major challenge in interpreting the results of hepatic vagal surgical denervation studies is that it interrupts both vagal sensory and parasympathetic cholinergic nerve fibers, which pass through the hepatic vagus nerve. Consequently, the specific role for parasympathetic cholinergic innervation of the liver plays in regulating hepatic lipid metabolism remain unclear.

Dynamic crosstalk between the liver and white adipose tissue (WAT) appears essential for hepatic fat accumulation and the development of MAFLD [9]. VLDL-TG synthesized in the liver is secreted and transported to adipose tissue for storage in the postprandial state [10].

Fasting increases lipolysis in adipocytes, and the free fatty acids released from WAT are transported to the liver for energy metabolism [10]. In human studies, most fatty acids constituting VLDL-TG are derived from adipose tissue [11]. However, in obesity, adipose tissue can no longer effectively store lipids, thus redirecting them toward other organs, primarily the liver [9]. Interestingly, the liver has the capability to control lipid uptake and lipolysis in WAT by activating bile acid (BA) signaling pathways [12]. Vagotomized rats exhibit reduced expression of the cholesterol 7 alpha-hydroxylase (*Cyp7a1*) gene, the rate-limiting enzyme in the classic bile acid synthesis pathway in the liver, while displaying increased bile acid levels in the serum and bile, in part via enhanced intestinal passive absorption of BAs [13]. Biliary epithelial cells in rats express the muscarinic acetylcholine receptor type 1, and its activation by the cholinergic agonist carbachol reduces bile flow [14]. These results suggest that the parasympathetic cholinergic innervation to the liver has the potential to regulate bile acid synthesis and outflow from the liver, which would subsequently result in a decrease in lipolysis within the WAT. In fact, a recent 3D/Airyscan imaging study has demonstrated vesicular acetylcholine transporter (vAChT)-positive parasympathetic nerve terminals on the bile duct in human liver tissues [15].

In this study, we aimed to investigate the impact of the loss of the parasympathetic brain-liver cholinergic axis on hepatic lipid metabolism in mice fed a high-fat diet (HFD). Our anterograde and retrograde neuronal mapping and 3D imaging of cleared liver tissues revealed that a subset of hepatocytes and cholangiocytes received vAChT-positive nerve terminals from the DMV. The removal of the liver-innervating parasympathetic cholinergic neurons prevented the development of hepatic steatosis. This decrease in hepatic fat accumulation was linked to the beiging of ingWAT caused by activation of the G protein-coupled bile acid receptor 1 (GPBAR1 or TGR5) in ingWAT.

## Results

### Parasympathetic cholinergic innervation of the liver

In our previous study [16], we showed that a subset of DMV cholinergic neurons projected to the liver. To extend and further confirm the presence of DMV cholinergic neurons innervating the liver, we injected a Cre-dependent viral tracer (adeno-associated virus (AAV).PHP.eB-FLEX-tdTomato [17]) into the liver of choline acetyltransferase (ChAT)-IRES-Cre (ChAT^Cre^) mice [18]. The ChAT^Cre^ transgenic strain has been used to selectively label parasympathetic cholinergic neurons [19]. At 3 weeks post-viral infection, we performed double immunocytochemistry with anti-ChAT and anti-tdTomato antibodies on the brain sections and found that the DMV displayed a small number of tdTomato-positive cells (Fig 1A). Importantly, all tdTomato-positive cells in the DMV were ChAT-expressing cells (n = 72 out of 72 neurons from 3 mice), and approximately 10 % of DMV cholinergic neurons expressed tdTomato (range, 7–14 %; n= 72 out of 660 cholinergic neurons from 3 mice).

**Fig 1.**
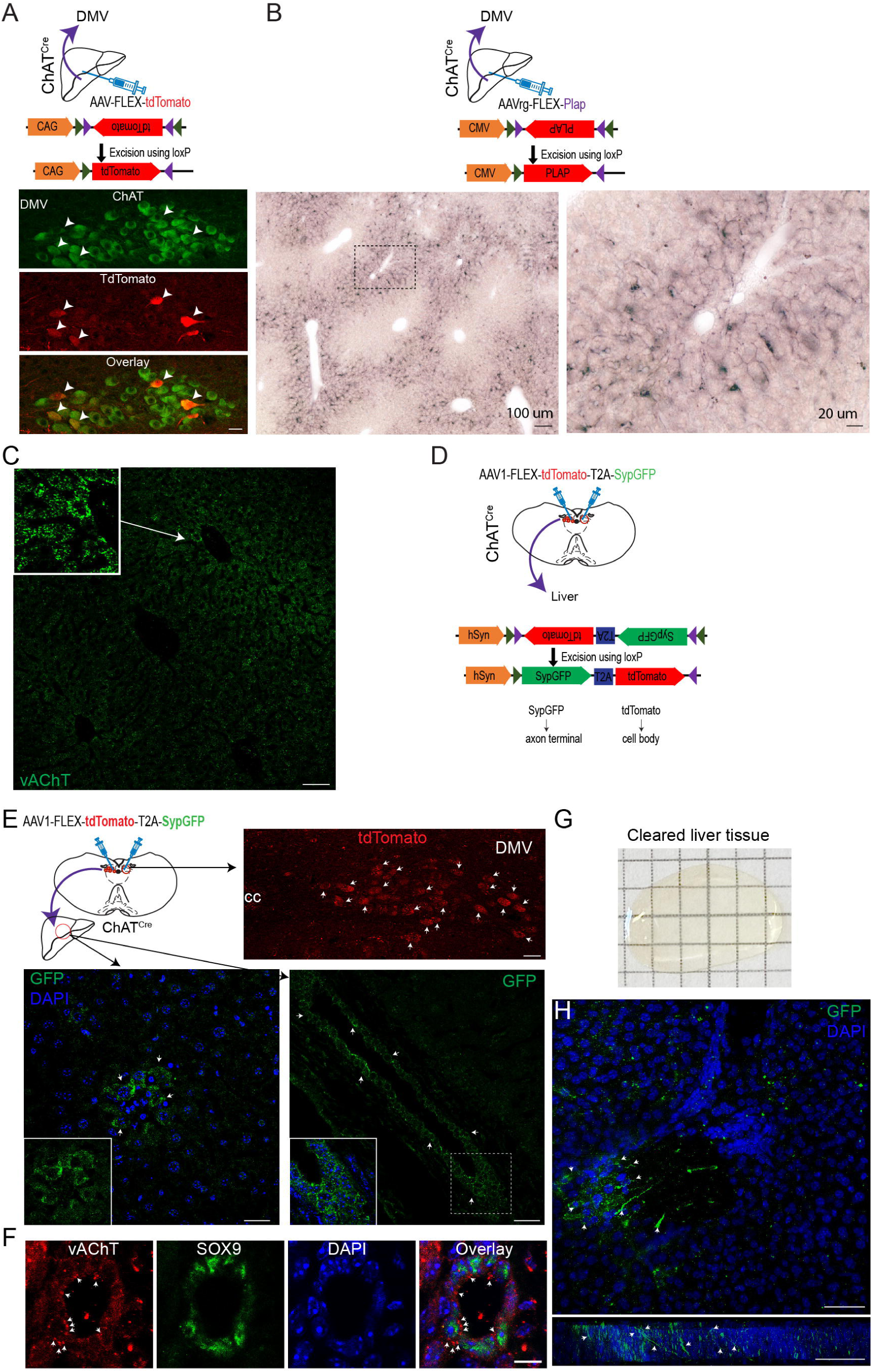
Parasympathetic cholinergic innervation of the mouse liver (A) Schematic illustration of our experimental conditions (Upper panel). Bottom panel: Images of confocal fluorescence microscopy showing co-expression of tdTomato and ChAT in a subset of DMV cholinergic neurons in ChAT^Cre^ mice receiving an injection of AAV.Php.eB-tdTomato into the liver, representing cholinergic neurons innervating the liver (white arrowheads). Scale bar, 30 μm (B) Schematic illustration of our experimental conditions (Upper panel). Bottom panel: Images showing PLAP-positive nerve fibers in the liver parenchyma. Right panel: Higher-magnification view of the black square box in the left panel. Scale bars, 100 μm (left), 20 μm (right) (C) Images of confocal fluorescence microscopy showing the expression of vAChT-positive nerve terminals (green) in the liver parenchyma (inset: higher-magnification view of vAChT-positive terminals). Scale bar, 70 μm (D) Schematic illustration of our experimental conditions. (E) Images of confocal fluorescence microscopy showing tdTomato-positive cholinergic neurons in the DMV (upper panel; white arrowheads) and sypGFP-positive nerve terminals (bottom panel; white arrowheads) in the liver of ChAT^Cre^ mice injected with AAV1-FLEX-tdTomato-T2A-sypGFP into the DMV. A small number of hepatocytes and cholangiocytes received parasympathetic cholinergic innervation (inset). Blue: nucleus staining with DAPI. Upper panel: scale bar, 30 μm; lower panel: left, scale bar, 20 μm; right, 40 μm (F) Images of confocal fluorescence microscopy showing that SOX9-positive cholangiocyte cells receive vAChT-positive nerve terminals (white arrowheads). Scale bar: 10 μm (G) Image of the liver after tissue clearing. (H) Z-stack image showing parasympathetic GFP-positive cholinergic nerve terminals (green) around the periportal area in cleared liver tissue collected from ChAT^Cre^ mice injected with AAV1-FLEX-tdTomato-T2A-sypGFP (upper panel). Bottom panel: 2D image of GFP-positive cholinergic nerve terminals shown in the upper panel. White arrowheads indicate the cholinergic nerve terminals. Scale bar: 50 μm

We subsequently examined whether these DMV cholinergic neurons send axonal projections to the liver by employing a retrograde AAV that encodes a Cre-dependent placental alkaline phosphatase (PLAP), which is expressed on the plasma membrane (AAVrg-FLEX- PLAP). Injection of this AAVrg-FLEX-PLAP into the liver of ChAT^Cre^ mice resulted in the presence of PLAP-positive nerves in the parenchyma of the liver, particularly in the periportal and midlobular areas (Fig. 1B). Immunostaining with an antibody against the presynaptic cholinergic marker vesicular acetylcholine transporter (vAChT) revealed the presence of vAChT- positive nerve terminals scattered throughout the liver parenchyma, but higher levels of vAChT- positive terminals were observed around the periportal zones (Fig 1C). In addition, the 3D projection view of the cleared liver tissue from the ChAT^Cre^;channelrodopsin (ChR2)-tdTomato mice showed the presence of hepatocytes receiving tdTomato-positive puncta-like nerve terminals (S1-Movie). Consequently, our findings align with prior studies indicating that a subset of DMV cholinergic neurons innervate the liver [16, 20, 21].

Given that DMV cholinergic neurons projected to the liver, we injected an anterograde AAV1 encoding Cre-dependent tdTomato and a synaptophysin-GFP (sypGFP) fusion protein separated by the sequence of a T2A self-cleaving peptide (AAV1-FLEX-tdTomato-T2A-sypGFP [22]) into the DMV of ChAT^Cre^ mice (Fig 1C). Under these experimental conditions, we anticipated that tdTomato would be expressed in DMV cholinergic neurons, and that sypGFP would move forward to the axon terminals of DMV cholinergic neurons. As shown in Fig. 1E, immunostaining revealed that the DMV cholinergic neurons were positive for tdTomato, indicating that the DMV cholinergic neurons were successfully infected. Immunolabeling of liver sections further revealed numerous sypGFP-positive nerve terminals in a subset of hepatocytes and bile duct epithelial cells cholangiocytes (Fig 1E and F). SRY-Related HMG Box Gene 9 (SOX9)-positive cholangiocytes were surrounded by vAChT-positive nerve terminals (Fig 1F).

3D imaging analysis of cleared liver tissue collected from ChAT^Cre^ mice injected with AAV1- FLEX-tdTomato-T2A-sypGFP further confirmed the presence of cholinergic nerve terminals around the portal vein and in a subset of hepatocytes (Fig 1G, H, and S2 Movie), consistent with a recent findings of parasympathetic cholinergic innervation of the bile duct in human liver tissues [15].

In addition to cholinergic input, RT-qPCR of liver tissue homogenates revealed the presence of muscarinic ACh receptor types 1, 2, 3, 4, and 5 (C*hrm*1, 2, 3, 4, and 5) (S1A Fig) consistent with our prior study [16]. Western blot analysis of liver tissue homogenates showed the expression of CHRM1, 2, 4 and 5, but not CHRM3 (S1B and C Fig). As CHRM4 protein expression was highly expressed compared to other CHRM subtypes, we immunolabeled the liver tissues with an anti-CHRM4 antibody. CHRM4 was detected in areas adjacent to vAChT-positive terminals (S1D Fig). Similarly, CHRM2 was also found in hepatocytes (S1E Fig). Taken together, our analyses of cleared liver tissues using 3D imaging techniques, as well as antero- and retrograde neural tracing robustly support the interpretation that the liver is innervated by parasympathetic cholinergic neurons.

### Liver-innervating parasympathetic cholinergic neurons are indispensable for the development of hepatic steatosis during a HFD feeding

The role of DMV cholinergic neurons innervating the liver in controlling hepatic lipid metabolism remains unexplored. Consequently, we examined the physiological impact of hepatic cholinergic loss-of-function on hepatic fat accumulation. First, to selectively target liver-projecting cholinergic neurons, we used a retrograde AAV (AAVrg) encoding a Cre-dependent diphtheria toxin subunit A fragment (dtA) that induces apoptosis of target cells [23]. AAVrg-FLEX-dtA and AAVrg-FLEX-tdTomato as control were injected into the livers of the ChAT^Cre^ mice (Fig 2A). To validate cholinergic cell death in the DMV, we sacrificed mice at 10 weeks post-viral inoculation and stained the DMV cholinergic neurons with an anti-ChAT antibody. The number of DMV cholinergic neurons was significantly lower in ChAT^Cre^ mice that received retroAAV-FLEX-dtA than in the controls (Fig 2B, C and S2A and B Fig). This reduced number of DMV cholinergic neurons was strongly associated with a lack of vAChT-positive nerve terminals in the livers of ChAT^Cre^ mice that received AAVrg-FLEX-dtA (Fig. 2D). Importantly, measurement of ACh levels revealed a significant reduction in hepatic ACh content in the experimental mice relative to controls, whereas there was no difference in plasma ACh levels between the experimental and control groups (Fig 2E and F). These results support the interpretation that our experimental approach effectively ablates parasympathetic cholinergic neurons innervating the liver.

**Fig 2.**
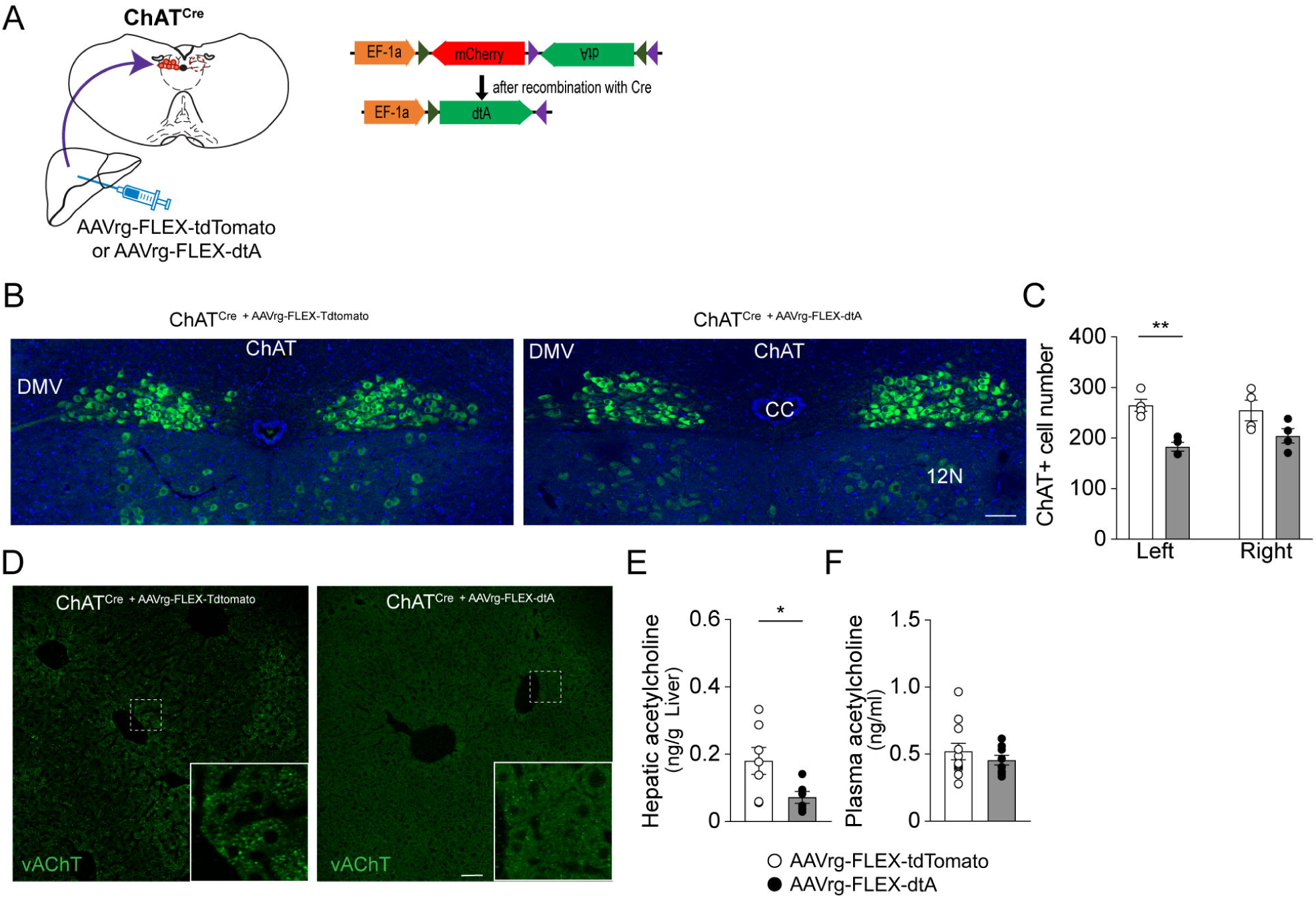
Loss of parasympathetic cholinergic neurons innervating the liver (A) Schematic illustration of the experimental configurations, where AAVrg-FLEX-tdTomato (control, ο) and AAVrg-FLEX-dTA (•) viral vectors were injected directly into the livers of ChAT^Cre^ mice. (B) Images of confocal fluorescence microscopy showing reduced numbers of cholinergic neurons in the DMV of ChAT^Cre^ mice receiving an injection of AAVrg-FLEX-dtA compared with the controls. Scale bar, 80 μm, cc: central canal, 12N: hypoglossal nucleus (C) Summary graph showing the number of cholinergic neurons on the left and right sides of the DMV in the control (n= 4 mice) and experimental mice (n= 4 mice). Unpaired *t*-test, **p<0.01 (D) Images showing a lack of vAChT-positive nerve terminals in the liver parenchyma of ChAT^Cre^ ^+ AAVrg-FLEX-dtA^ mice (right panel). Left panel: control. Scale bar, 50 μm (E and F) Summary plots showing a significant decrease in hepatic (control, n= 7 mice; experimental mice, n= 6 mice), but not plasma (control, n= 11 mice; experimental mice, n= 8 mice), ACh contents in ChAT^Cre^ ^+ AAVrg-FLEX-dtA^ mice compared to the controls. Unpaired *t*-test, *p<0.05 The data supporting the graphs shown in the figure (Fig. 2C,E,F) are available in the S1 Data file.

We then examined whether deleting liver-innervating cholinergic neurons alters hepatic lipid metabolism during HFD feeding. To address this question, we first investigated if there are differences in the liver histology between the groups. Liver tissues were labeled with liver zonation markers such as the pericentral marker glutamine synthetase (GS) and the periportal marker E-cadherin (E-cad) [24]. The expression pattern of these zonation markers in the livers of the control group fed a standard chow diet demonstrated that GS- and E-cad-positive areas were clearly separated in the liver (S3A Fig). Interestingly, high-fat feeding robustly reduced E- cad expression in periportal hepatocytes in the livers of control mice compared to control mice fed a standard chow diet (Fig 3A), consistent with disrupted E-cad expression in chronic liver disease in mice [25]. Surprisingly, deleting the parasympathetic cholinergic input to the liver in ChAT^Cre^ mice fed HFD restored the expression of E-cad in periportal hepatocytes (Fig 3B). In contrast, GS was similarly expressed in pericentral hepatocytes in both control and experimental mice fed HFD (Fig 3A and B).

**Fig 3.**
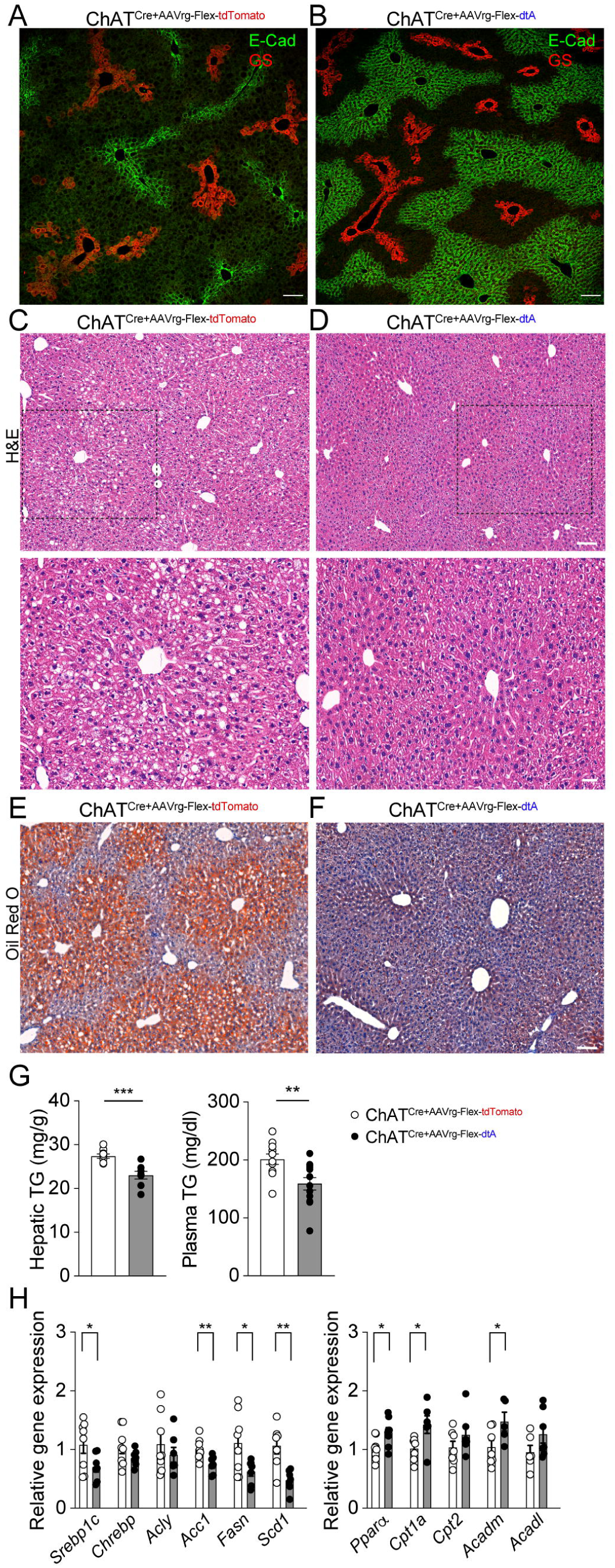
Deleting liver-innervating cholinergic neurons prevents hepatic steatosis during high-fat diet feeding. (A and B) Macroscopic appearance of livers of ChAT^Cre^ mice receiving AAVrg-FLEX-tdTomato and AAVrg-FLEX-dtA. E-Cad expression was lower in the control group than in the experimental group. Scale bar, 100 μm (C and D) H & E staining of liver tissues revealed increased numbers of lipid droplets and ballooned hepatocytes in the controls compared to those in the experimental mice (top panel). Scale bar, 100 mm. Bottom panel: Higher magnification view of liver tissue in the black dotted square. Scale bar, 30 μm (E and F) Oil red O staining (red) of liver tissues from ChAT^Cre^ mice with and without hepatic cholinergic input. The experimental group did not exhibit fat accumulation in the liver parenchyma during high fat feeding. Nuclear staining (blue): hematoxylin, Scale bar, 100 μm (G) Graphs showing hepatic (n= 9 vs. 8 mice) and plasma (n= 11 vs. 12 mice) TG levels in the controls and experimental mice. There was a significant reduction in both hepatic and plasma TG levels in the experimental group relative to those in the control group (unpaired *t*-test, **p<0.01, ***p<0.001). (H) Summary graphs showing changes in hepatic mRNA expression of enzymes involved in lipid metabolism. *Srebp1*c mRNA expression was significantly downregulated, while *Ppara* mRNA expression was increased in the livers of ChAT^Cre^ mice receiving AAVrg-FLEX-dtA relative to controls (control, n= 6-8 mice; experimental, n= 6-8 mice, unpaired t-test, *p<0.05, **p<0.01). The data supporting the graphs shown in the figure (Fig. 3G,H) are available in the S1 Data file.

Hematoxylin and eosin (H & E) staining revealed a clear difference in the macroscopic appearance of the livers of ChAT^Cre^ mice receiving AAVrg-FLEX-tdTomato and AAVrg-FLEX-dtA. Control mice exhibited an increased number of large and small lipid droplets and ballooned hepatocytes compared to experimental mice (Fig 3C and D). Oil Red O staining further showed that ChAT^Cre^ mice without DMV cholinergic neurons innervating the liver contained fewer lipid droplets in the liver than controls (Fig 3E and F). These results suggest that parasympathetic cholinergic input to the liver limits hepatic steatosis during high-fat diet feeding.

As there was a lack of lipid droplets in the livers of the experimental mice, we measured the hepatic and plasma triglyceride (TG) levels. Following HFD feeding, ChAT^Cre^ mice without parasympathetic cholinergic neurons innervating the liver on high-fat feeding exhibited reduced hepatic and plasma TG levels in the liver relative to controls (Fig 3G). This was associated with decreased plasma TG levels (Fig 3G). Reverse transcription-quantitative polymerase chain reaction (RT-qPCR) analysis of liver tissues revealed a significant decrease in the expression of the lipogenic transcription factor sterol regulatory element-binding protein 1-c (*Srebp-1*) in the experimental mice relative to the controls. The experimental mice also exhibited downregulation of its target genes, including acetyl-coenzyme A carboxylase 1 (*Acaca*), fatty acid synthase (*Fasn*), stearoyl-CoA desaturase-1 (*Scd1*) (Fig 3H). In addition, the ChAT^Cre^ mice without parasympathetic cholinergic input to the liver had increased expression of peroxisome proliferator-activated receptor alpha (PPAR-α), the central transcriptional regulator of fatty acid oxidation relative the control group. As a result, there was increased carnitine palmitoyltransferase 1a (*Cpt1a*) and acyl-CoA dehydrogenase medium chain (*Acadm*) mRNA expression in the experimental mice relative to controls (Fig 3H). In contrast to male mice, high-fat feeding did not cause any changes in liver histology or fat accumulation in both control and experimental females (S4A-C Fig). In fact, hepatic steatosis was less common in C57BL/6 females than in males during high-fat feeding [26–28].

Hepatic vagal denervation results in an increase in hepatic interleukin 6 (*Il6*) gene expression in mice, which subsequently modulates the expression of hepatic gluconeogenic enzymes [29]. In our experiments, where we selectively deleted liver-projecting cholinergic neurons, we observed a significant decrease in Il6 gene expression (S5A Fig). Notably, there was no significant difference in gluconeogenic gene expression between the experimental and control groups (S5B. Fig). Furthermore, we investigated whether the loss of parasympathetic cholinergic innervation affects sympathetic input to the liver. We used immunostaining with an anti-tyrosine hydroxylase (Th) antibody and found no detectable difference in Th-positive nerve fibers (S6A. and B. Fig). Additionally, hepatic norepinephrine levels were similar between the groups (S6C. Fig). Taken together, these findings suggest that the loss of parasympathetic cholinergic neurons innervating the liver prevents the development of hepatic steatosis during HFD feeding in male mice.

The male control mice displayed dense lipid accumulation in the pericentral areas compared to the experimental mice. Therefore, we further examined if there are distinct gene expression patterns throughout the hepatic zonation in both groups by performing spatial transcriptomics (S7. Fig). Pericentral, midlobular, and periportal clusters were identified based on known zonation markers (S7A. Fig). We found that the midlobular areas exhibited 391 differentially expressed genes (DEGs) (S7B. Fig and S1. Table) and that 271 genes in the PC and 550 genes in the PP areas were significantly different between the groups (S7B. Fig and S2 and 3. Table). Importantly, the loss of liver-projecting cholinergic neurons downregulated hepatic lipogenesis-related genes, including *Scd1*, *Fabp4*, *Cd36*, *Lpin1*, *Nampt*, *Nr1l2*, and *Me1* and upregulated the *Trib1* gene across the liver zonation (S7C. Fig).

A selective gene ontology (GO) analysis of the DEGs in each zonation using Enrichr-KG [30] identified the top 5 enriched GO terms in biological process (S7D. Fig). These included "cotranslational protein targeting to membrane" (GO:0006613), "SRP-dependent cotranslational protein targeting to membrane" (GO:0006614), "protein targeting to ER" (GO:0045047), "cytoplasmic translation" (GO:0002181), and "cellular protein metabolic process" (GO:0044267). According to the mammalian phenotype ontology analysis [31], each zonation shared 3 terms, including "abnormal lipid homeostasis" (MP:0002118), "abnormal liver physiology" (MP:0000609), and "hepatic steatosis" (MP:0002628). These results further support that liver-projecting cholinergic neurons play an important role on developing hepatic steatosis in mice fed HFD.

### Loss of liver-innervating cholinergic neurons causes beiging of ingWAT

One of the hallmarks of diet-induced obeisty is a set of increases in inguinal WAT depot size, accompanied by increased adipocyte cell size [32]. Importantly, we found that ChAT^Cre^ mice without liver-innervating cholinergic neurons had reduced ingWAT size relative to control mice (Fig. 4A). Histological analysis of ingWAT revealed smaller adipocyes and decrease adipocyte area in experimental mice compared to the control mice (Fig 4B, C, and D). These results suggest that a loss of DMV cholinergic input to the liver prevents ingWAT hypertrophy.

**Fig 4.**
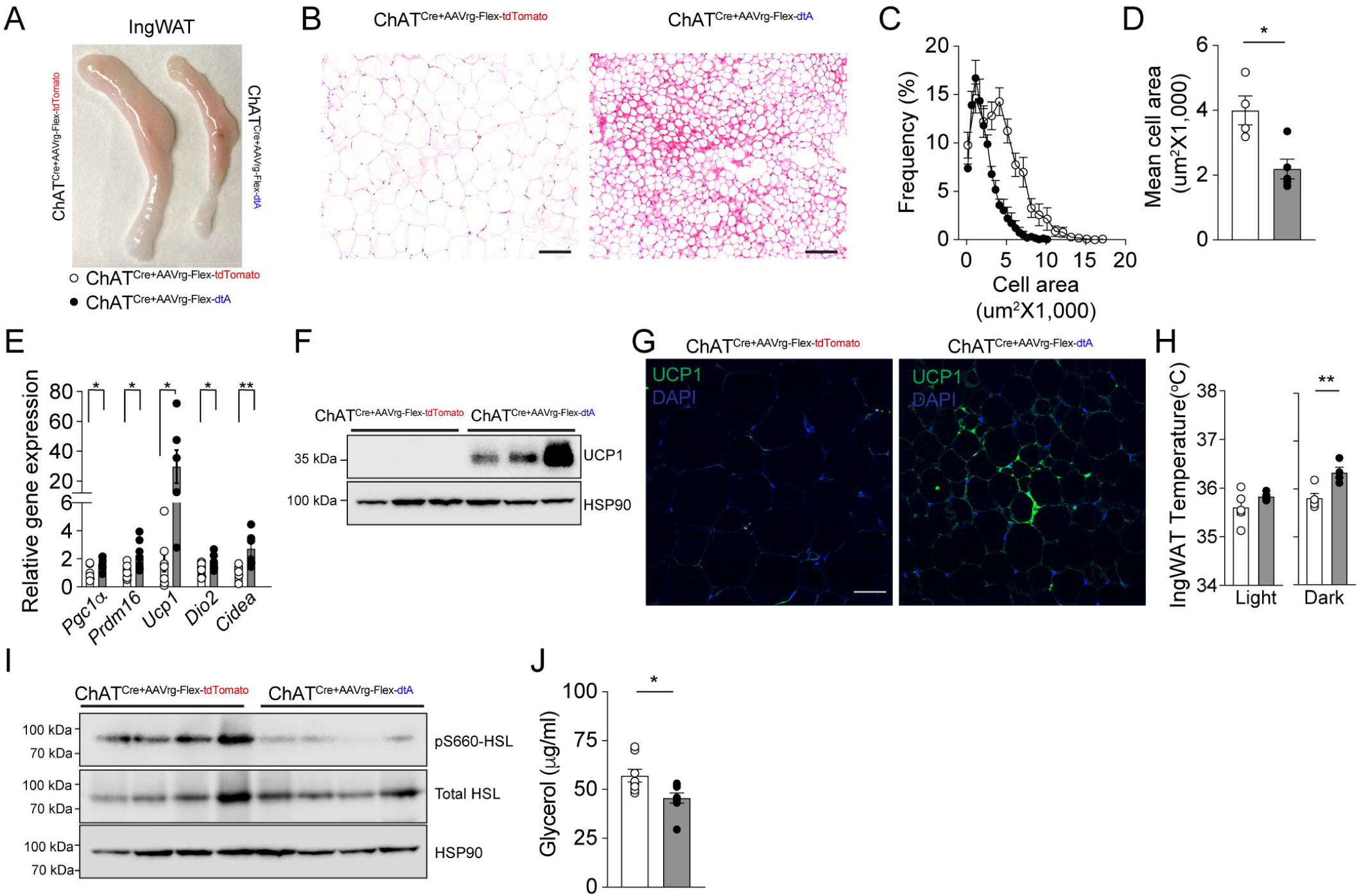
Deleting liver-innervating cholinergic neurons induces beiging of ingWAT (A) Representative images of the ingWAT of control and experimental mice. (B) Representative images of adipocytes in the ingWAT of control and experimental groups. Scale bar, 100 μm. (C and D) Plots showing the distribution of the average adipocyte size (left panel). Right panel: The mean cell area was significantly smaller in the experimental groups than in controls (control, n=4 mice; experimental, n=5 mice, unpaired *t*-test, *p<0.05) (E) Summary graph showing changes in thermogenic gene expression in ingWAT of ChAT^Cre^ mice with and without DMV cholinergic neurons innervating the liver. Ablation of liver-projecting cholinergic neurons significantly upregulated the thermogenic gene expression (unpaired *t*-test, *p<0.05, **p<0.01). (F and G) Western blot images of ingWAT showing an increase in UCP1 expression in experimental mice. Images of confocal fluorescence microscopy showing increased UCP1 expression in the ingWAT of experimental mice (G). Scale bar, 50 μm. (H) Summary plots showing ingWAT temperature in the light and dark phases (control, n=5 mice; experimental, n= 4 mice, unpaired *t*-test, **p<0.01) (I) Western blot images showing a decrease in pS660-HSL expression in ingWAT of experimental mice relative to controls. (J) Summary plot showing plasma glycerol levels in the control (n= 8 mice) and experimental mice (n= 8 mice, unpaired *t*-test, *p<0.05) The data supporting the graphs shown in the figure (Fig. 4C, D, E, H and J) are available in the S1 Data file.

RT-qPCR analysis revealed that deleting DMV cholinergic neurons innervating the liver significantly upregulated the expression of Peroxisome proliferator-activated receptor-gamma coactivator (PGC)-1alpha (PGC1a) in ingWAT, which is required for the expression of mitochondrial biogenesis and thermogenic genes in brown fat cells [33, 34] (Fig 4E). Likewise, the expression of PR (PRD1-BF1-RIZ1 homologous)-domain containing 16 (PRDM16), a transcription factor that plays a key role in browning and thermogenesis of subcutaneous WAT [35], was significantly increased in experimental mice relative to controls (Fig 4E). The upregulation of these genes was associated with a robust increase in the mRNA expression of uncoupling protein 1 (*Ucp1*), iodothyronine deiodinase 2 (*dio2*), and cell death-inducing DFFA-like effector A (*Cidea*) (Fig 4E). Western blotting and immunostaining further showed an increase in UCP1 protein levels in ingWAT of ChAT^Cre^ mice receiving AAVrg-FLEX-dtA to the liver (Fig 4F and G). In addition, daily measurement of ingWAT temperature revealed a significant elevation in the ingWAT temperature of the experimental mice in the dark phase (Fig 4H). These data support that a loss of parasympathetic cholinergic neurons innervating the liver induces beiging of ingWAT, resulting in an increase in thermogenesis.

WAT lipolysis in obesity can result in an excess of fatty acids and glycerol in the liver [36]. This lipid influx can cause hepatic TG synthesis, which results in MAFLD [37]. We investigated whether eliminating cholinergic input to the liver from the DMV would decrease HSL-mediated lipolysis in ingWAT. The Western blot analysis of ingWAT demonstrated a decrease in the levels of phospho-HSL in the experimental mice (Fig 4I). Moreover, we observed a decrease in plasma glycerol levels in the experimental mice compared to the control mice (Fig 4J). Taken together, these findings demonstrate that the loss of hepatic cholinergic innervation led to an increase in the expression of thermogenic genes while simultaneously reducing lipolysis in ingWAT.

### Increased bile acid secretion induces ingWAT beiging via TGR5

How would removing hepatic cholinergic innervation drive these reductions in adiposity in the ingWAT, which does not receive parasympathetic cholinergic innervation in rodents [38, 39]. Our results suggest that removing hepatic parasympathetic cholinergic input enhances liver-adipose crosstalk, limiting ingWAT lipolysis and thereby limiting hepatic steatosis during HFD feeding. Previous studies have indicated that bile acid and its corresponding receptor TGR5 are crucial in the browning of WAT [12, 40]. We thus investigated whether the loss of function of liver-innervating cholinergic neurons affects the expression of genes related to bile acid synthesis, such as *Cyp7a1* and *Cyp7b1*. We found that hepatic cholinergic loss of function upregulated expression of the *Cyp7b1* gene involved in an alternative bile acid synthesis pathway [41] (Fig 5A). Furthermore, our spatial transcriptomic analysis of the liver revealed that ChAT^Cre^ mice without DMV cholinergic neurons innervating the liver displayed a trend toward an increase in *Cyp7b1* gene expression in all three zones (Fig 5B), whereas *Cyp7a1* expression was not altered, as observed in qPCR experiments (Fig 5A and 5B). Elevated *Cpy7b1* expression led to a substantial increase in fasting serum bile acid levels in ChAT^Cre^ mice that received AAVrg-FLEX-dtA. (Fig 5C). In our further analysis of the fasting serum bile acid profiles in both control and experimental mice, we observed a significant increase in primary bile acids such as cholate and chenodeoxycholate, which was attributed to the loss of hepatic cholinergic input (Fig 5D).

**Fig 5.**
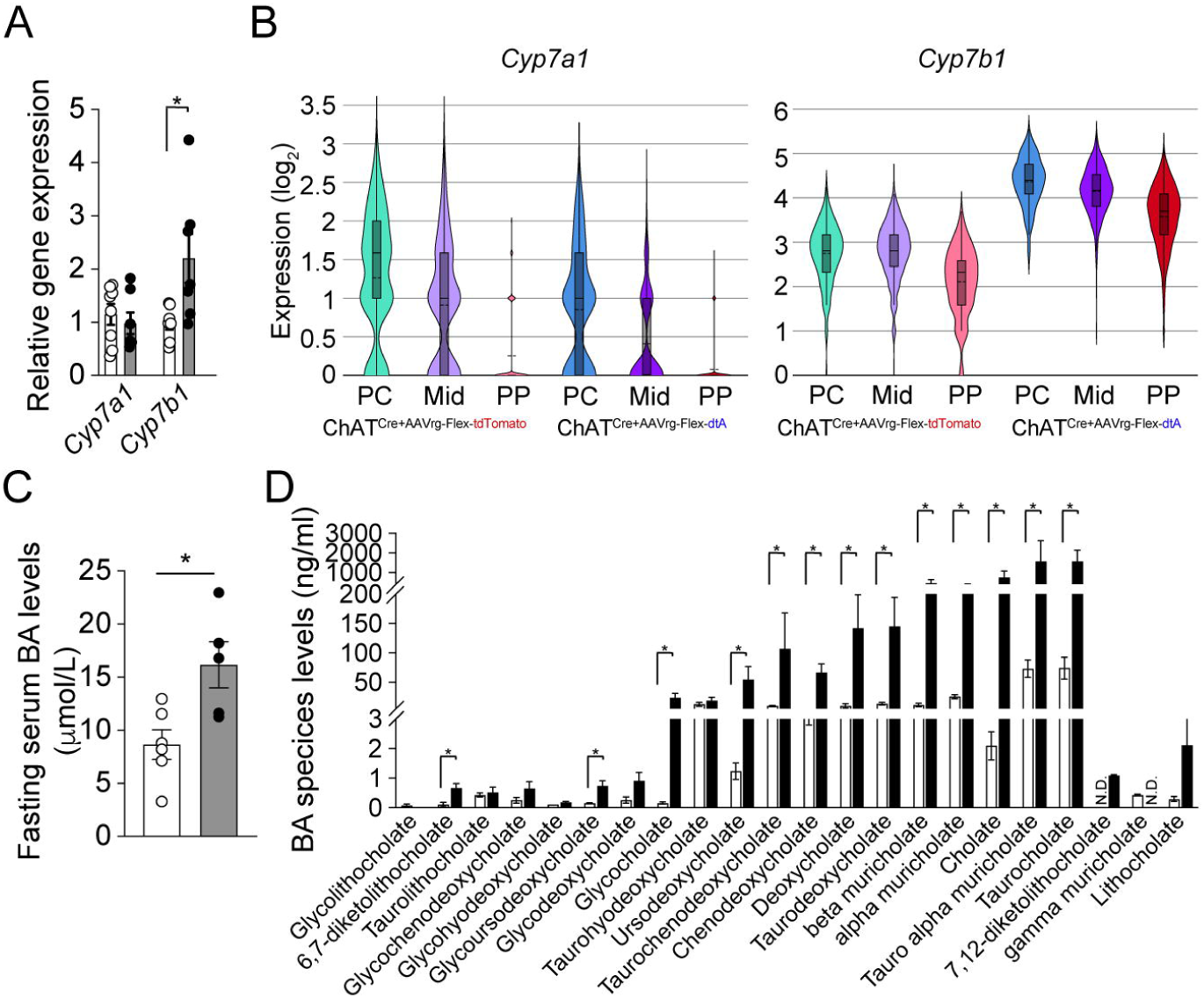
Ablation of liver-innervating cholinergic neurons elevates serum bile acid levels. (A) Graph showing mRNA expression of the enzymes involved in bile acid synthesis in liver. While there was no difference in *Cyp7a1* gene expression, the loss of hepatic cholinergic input significantly upregulated *Cyp7b1* mRNA expression (control (open circle), n=7-8 mice; experimental (closed black circle), n=7 mice). (B) Graphs showing *Cyp7a1* and *Cyp7b1* gene expression in the periportal (PP), midlobular (Mid), and pericentral (PC) zones in the livers of the controls and the experimental groups. (C) Graph showing the fasting serum bile acid levels in the control group (n=6 mice) and the experimental group (n=5 mice). Unpaired t-test, *p<0.05. (D) Graph showing fasting serum bile acid profile in the control (n=4 mice) and experimental (n= 4 mice) mice. The Mann-Whitney test revealed a statistically significant difference between the two groups (*p<0.05). The data supporting the graphs shown in the figure (Fig. 5A-D) are available in the S1 Data file.

We then examined whether the expression of its corresponding receptor, TGR5, is altered in the ingWAT of the experimental mice. Our observations revealed elevated levels of both TGR5 gene and protein expression in the ingWAT of the experimental mice compared to the control group (Fig 6A and B). This suggests that the increase in serum bile acids and *Tgr5* expression in the ingWAT leads to beiging of the ingWAT in the experimental mice. To investigate this possibility, we administered *Tgr5* siRNA or control siRNA into the ingWAT of ChAT^Cre^ mice without liver-innervating cholinergic neurons to exclusively knock down *Tgr5* in the ingWAT (Fig 6C). Ten weeks after a viral infection, we conducted RT-qPCR and Western blot analyses on samples and found that the *Tgr5* gene and protein levels in ingWAT were significantly reduced following the injection of *Tgr5* siRNA (Fig 6D), demonstrating the effectiveness of our approach in knocking down *Tgr5* expression. Our experiments also revealed that the number of large-sized adipocytes and crown-like structures in the ingWAT of mice that received *Tgr5* siRNA was increased (Fig 6E), accompanied by a notable increase in adipocyte area (Fig 6F and G).

**Fig 6.**
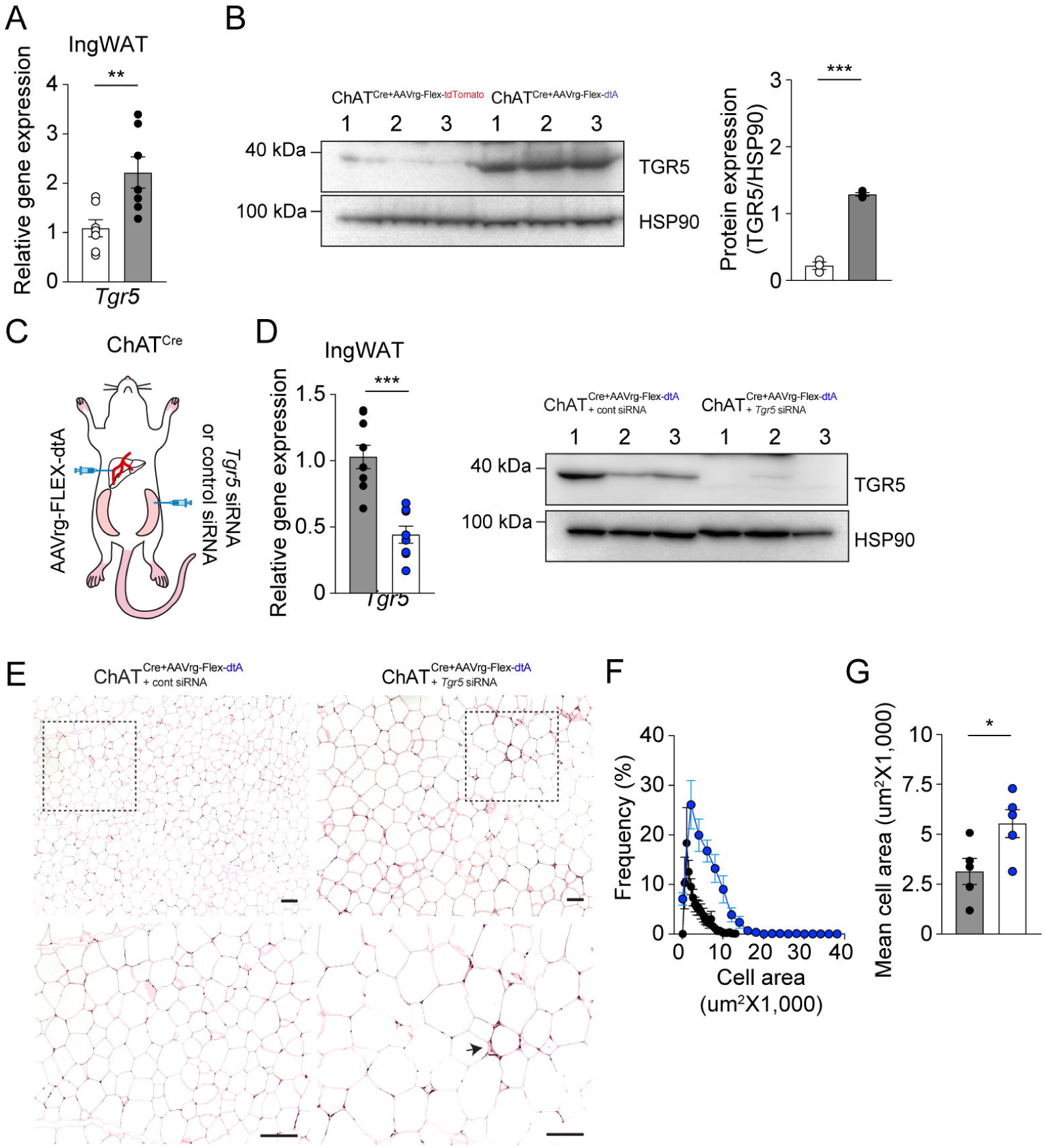
Liver-ingWAT communication contributes to beiging of ingWAT via bile acids. (A and B) Graphs showing mRNA and protein expression levels of *Tgr5* in ingWAT. There was a significant increase in TGR5 expression in the ingWAT of ChAT^Cre^ mice that received AAVrg-FLEX-dtA (RT-qPCR, control (open circle), n=7 mice; experimental (closed circle), n=7 mice, unpaired *t*-test, **p<0.01; Western blotting, control (open circle), n=3 mice; experimental (closed circle), n=3 mice, unpaired *t*-test, ***p<0.001). (C) Schematic illustration showing our experimental configurations. (D) Reduction in *Tgr5* gene and protein levels in ChAT^Cre^ mice following administration of AAVrg-FLEX-dtA to the liver (black circle) and another injection of *Tgr5* siRNA into both sides of the ingWAT (blue circle)(unpaired *t*-test, *** P <0.001). (E) Representative images of ingWAT sections stained with H&E. ChAT^Cre^ mice without DMV cholinergic neurons innervating the liver receiving an injection of *Tgr5* siRNA into ingWAT exhibited larger adipocytes compared to the control mice (upper panel). Scale bar, 50 μm Bottom panel: higher magnification view of the black dotted square. Arrow represents crown-like structure. Scale bar, 100 μm (F and G) Plot showing the distribution of average adipocyte size. The mean cell area was significantly larger in the experimental group receiving *Tgr5* siRNA than in the experimental group receiving control siRNA (I) (control, n=5 mice; experimental, n=5 mice, unpaired *t*-test, *p<0.05) The data supporting the graphs shown in the figure (Fig. 6A,B,D,F and G) are available in the S1 Data file.

We examined whether *Tgr5* knockdown impacts HSL-mediated lipolysis in ingWAT. Our findings showed an increase in phospho-HSL levels in the ingWAT (Fig 7A), which subsequently led to higher levels of plasma glycerol (Fig 7B). Considering the well-established connection between elevated plasma glycerol levels and lipid accumulation in the liver, we sought to determine if *Tgr5* knockdown results in hepatic steatosis. Our results revealed the existence of large lipid droplets and ballooned hepatocytes in the liver sections of mice that received *Tgr5* siRNA (Fig 7C). Furthermore, Oil Red O staining confirmed the accumulation of lipids in the livers of mice treated with *Tgr5* siRNA in the ingWAT (Fig 7D). Additionally, we found that the knockdown of the Tgr5 gene in the ingWAT of ChAT^Cre^ mice without DMV cholinergic neurons to the liver resulted in a significant increase in hepatic TG content, which is indicative of enhanced lipid accumulation in the liver (Fig 7E). Collectively, our findings support the idea that cross-talk between the liver and ingWAT through elevated bile acid signaling plays a vital role in maintaining the lipid metabolism balance in both the liver and ingWAT.

**Fig 7.**
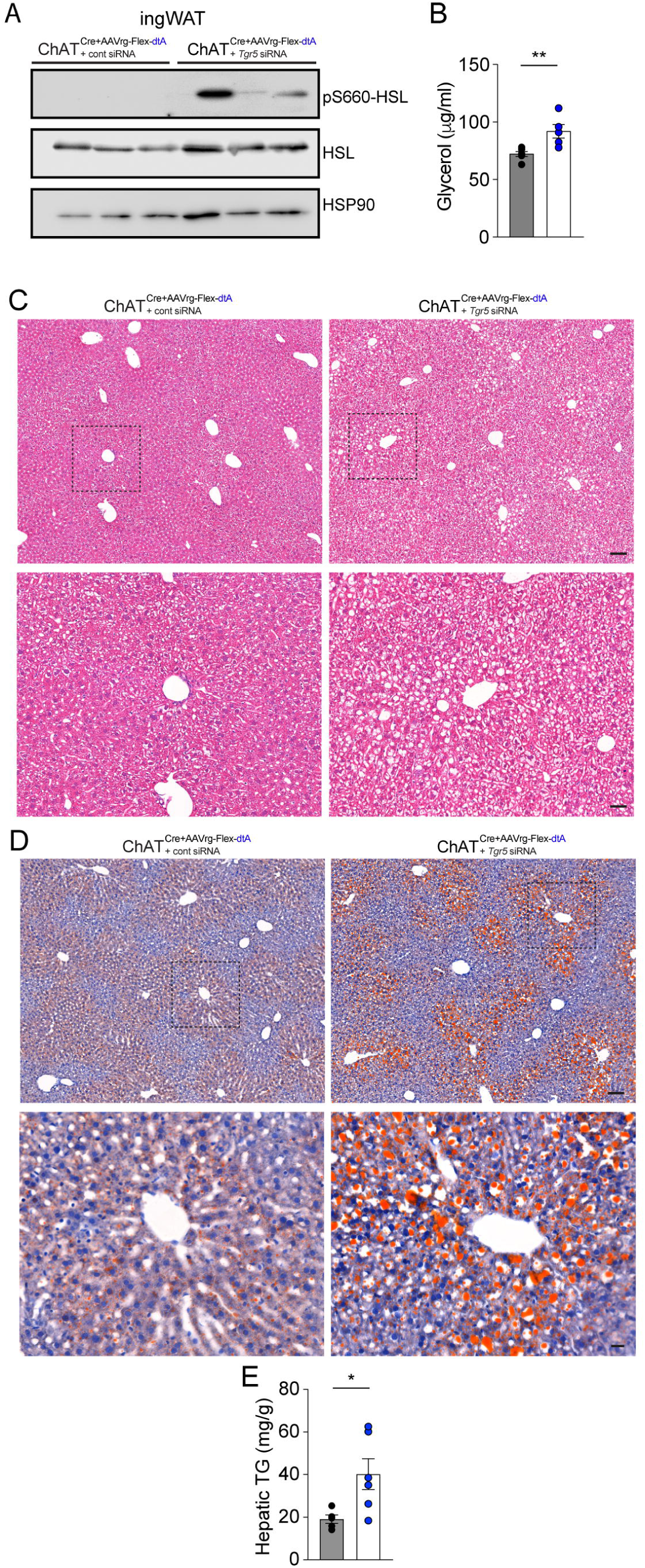
*Tgr5* knockdown in ingWAT causes hepatic steatosis in ChAT^Cre^ mice receiving AAVrg-FLEX-dtA during high-fat diet feeding. (A) Gel images showing the Western blot analysis of ingWAT from experimental mice revealed an increase in the expression of pS660-HSL when compared to the control group. (B) Graph showing plasma glycerol levels in the controls (black circle, n= 6 mice) and the experimental groups (blue circle, n= 5 mice). Unpaired *t*-test, **p<0.01 (C and D) Representative images of liver sections stained with H&E (C) and Oil Red O (D). Upper panel: scale bar, 100 μm; bottom panel: higher magnification view of the black dotted square scale bar, 50 μm (E) Graph showing hepatic TG levels in the controls (n= 5 mice) and the experimental groups (n= 6 mice) . Unpaired *t*-test, *p<0.05 The data supporting the graphs shown in the figure (Fig. 7B and E) are available in the S1 Data file.

### Deleting the brain-liver axis has a slight but significant effect on body weight

We also investigated whether the disruption of the brain-liver axis could impact body weight in mice fed HFD. After 10 weeks of high-fat diet feeding, we observed a small but significant difference in body weight gain between the control and experimental groups (S8A Fig). The experimental mice gained less weight than the control mice during the HFD period. This decrease in weight gain was closely related to a significant reduction in fat mass in the experimental group, while lean mass was similar between the two groups (S8B Fig). When we compared adipose tissue mass collected from controls with that of adipose tissue from the experimental group, we found a significant decrease in the mass of ingWAT (S8C Fig). It is important to note that both groups consumed an equal amount of food during the light and dark phases (S8D Fig).

The reduction in body weight, despite no change in energy intake, may be due to an increase in energy expenditure. To assess metabolic rate, we placed animals in metabolic cages and observed that the experimental mice consumed more oxygen and produced more carbon dioxide during the dark phase than the controls (S8E and F Fig). These findings indicate an increase in total energy expenditure in the experimental mice (S8G Fig). However, the respiratory exchange ratio was similar between the two groups (S8H Fig). Notably, the experimental mice exhibited more locomotor activity during the dark phase (S8I and J Fig). Collectively, these results suggest that deleting liver-projecting cholinergic neurons plays a role in increasing energy expenditure.

### Loss of the brain-liver axis improves insulin tolerance in male mice fed HFD

We evaluated whether deleting the brain-liver axis affects glucose homeostasis in mice fed a high-fat diet, as the liver plays a crucial role in regulating systemic glucose levels [42]. To assess glucose intolerance and insulin sensitivity, we measured blood glucose and insulin levels. Our results showed that basal glucose levels were lower in ChAT^Cre^ mice receiving AAVrg-FLEX-dtA injection than in the controls (S9A Fig). Additionally, the loss of DMV cholinergic neurons innervating the liver led to a significant reduction in fasting glucose levels (S9B Fig).

To evaluate whether ChAT^Cre^ mice without parasympathetic cholinergic neurons innervating the liver exhibit any changes in their ability to dispose a glucose load, we conducted intraperitoneal glucose tolerance tests (i.p.GTT) and found no significant difference in blood glucose levels in response to a bolus of glucose (i.p., 2 g/kg) between the groups (S9C and D Fig). However, the experimental mice showed increased insulin sensitivity (S9E Fig). During the insulin tolerance tests (ITT) (i.p., 1 U/kg), blood glucose levels were significantly lower in ChAT^Cre^ mice injected with AAVrg-FLEX-dtA than in controls (S9E Fig). Furthermore, the improved insulin tolerance in the experimental group was associated with reduced plasma insulin levels relative to those in the controls (S9F Fig). These results indicate that the loss of DMV cholinergic input to the liver enhanced insulin sensitivity in ChAT^Cre^ male mice fed HFD.

Intriguingly, a growing number of studies have revealed sex-specific differences in metabolic homeostasis, diabetes, and obesity in both rodents and humans [43–49]. For instance, premenopausal women were relatively protected from diseases associated with metabolic syndrome [50]. Female rodents were less prone to developing diet-induced obesity (DIO), insulin resistance, hyperglycemia, and hypertriglyceridemia [43, 44, 47, 48]. In particular, C57BL/6 female mice were less likely to develop DIO and hyperglycemia than C57BL/6 male mice during high-fat feeding [47–49, 51, 52]. Our study also found that control female mice did not develop DIO or hyperglycemia (S10A-F Fig), and hepatic cholinergic loss of function had no effect on body weight or glucose homeostasis (S10A-F Fig). In other words, both groups did not show any notable differences in body weight, food intake, body composition, blood glucose levels, GTT, and ITT compared with the controls (S10A-H Fig).

## Discussion

The current study provides concrete evidence for the existence of the cholinergic system in the mouse liver. Our study showed that DMV cholinergic neurons directly projected to hepatocytes and cholangiocytes and that hepatocytes expressed muscarinic ACh receptors.

Our 3D imaging analysis of cleared liver tissue also revealed the presence of cholinergic nerve terminals around the portal vein. Additionally, our study found that deleting cholinergic neurons innervating the liver prevented the development of hepatic steatosis during HFD feeding, which was closely linked to the browning of ingWAT. ChAT^Cre^ mice without cholinergic input to the liver displayed elevated bile acid synthesis, primarily through the CYP7B1-mediated alternative pathway, leading to elevated serum bile acid levels. Importantly, the experimental mice exhibited elevated *Tgr5* gene expression in ingWAT, which was found to be critical in mediating the beneficial effects of the loss of hepatic cholinergic function. Specifically, knockdown of the *Tgr5* gene in ingWAT completely abolished these beneficial effects and led to the reappearance of hepatic steatosis in the experimental groups. These findings suggest that TGR5 is a crucial mediator of the metabolic benefits conferred by loss of hepatic cholinergic function. Taken together, our results suggest that the brain-liver axis plays an essential role in bile acid synthesis and secretion. The interorgan crosstalk via bile acids is also critical for the development of hepatic steatosis.

The liver plays a major role in controlling energy metabolism, including macronutrient metabolism, lipid homeostasis, and energy storage [10]. Like other visceral organs [53–55], the liver is innervated by the vagus nerve [56], and vagal innervation to the liver regulates these metabolic functions [5, 8, 57, 58]. Parasympathetic (or vagal) efferent neurons innervating the liver are located in the DMV [59]. Prior anatomical studies with retrograde neuronal tracers such as cholera toxin B, pseudorabies virus, and AAV encoding a Cre-dependent reporter protein demonstrated that the mouse liver receives parasympathetic nerve innervation from the DMV [16, 20, 21]. DMV motor neurons are acetylcholine (ACh)-expressing neurons [53]. We recently showed that hepatocytes received direct DMV cholinergic input and expressed muscarinic ACh receptor genes [16]. Importantly, a short-term change in the activity of DMV cholinergic neurons projecting to the liver regulated hepatic glucose output in lean mice [16]. In contrast to these prior findings, recent 3D imaging studies revealed a paucity of parasympathetic cholinergic innervation in the mouse liver [4, 60], indicating that hepatic metabolism is primarily regulated by the sympathetic nervous system. However, a recent elegant study by the Tang research group clearly demonstrated the presence of vAChT-positive nerve terminals in the bile duct of human liver tissues, suggesting that the liver receive parasympathetic cholinergic innervation [15].

In fact, accumulated studies have shown that parasympathetic innervation to the liver controls hepatic glucose output, lipid metabolism, and bile formation [5, 7, 8, 57, 61]. However, there is limited information available on the detailed neuroanatomy of parasympathetic innervation of the liver, leading some to question its importance in controlling hepatic metabolism. [4, 60, 62]. In addition, although hepatic branch vagotomy altered multiple physiological processes, including glucose homeostasis [57, 61], hepatic lipid metabolism [5, 7], and hepatic inflammatory response [8], studies involving hepatic branch vagotomy have not provided conclusive evidence supporting the role of the parasympathetic cholinergic system in liver function. The hepatic branches of the vagus nerve also contained sensory nerves that transmit interoceptive information from the liver to the brain. Despite this, several studies have used virus-mediated tracing methods to clearly demonstrate the presence of DMV cholinergic neurons that innervate the liver [16, 20, 21].

Our study, which utilized anterograde tracing techniques alongside 2D immunostaining and 3D imaging of cleared liver tissue, uncovered direct synaptic connections between cholinergic neurons in the DMV and the liver. The elimination of DMV cholinergic neurons that innervate the liver led to a substantial reduction in the number of vAChT-positive nerve terminals and hepatic ACh levels. It is possible that differences in the morphology of preganglionic and postganglionic neurons may contribute to the discrepancy between our findings and recent 3D imaging studies [4, 60]. While sympathetic postganglionic neurons and sensory neurons, particularly c-fibers, have unmyelinated nerve fibers, the nerve fibers of parasympathetic cholinergic preganglionic neurons are myelinated [63, 64]. This myelin sheath may make it difficult to stain cholinergic nerve fibers in the liver parenchyma. In addition, it is important to emphasize that the cholinergic nerves form a complex plexus with multiple axonal varicosities (en-passant synapses) in both the central and peripheral nervous systems [65–68]. Terminal synapses are typically established at the end of an axon’s projections, while en passant synapses are situated along the axon shaft, often at a considerable distance from the terminal. All these prior studies using the vAChT antibody showed that staining with the vAChT antibody consistently revealed puncta-like varicosities. To circumvent this issue, we expressed PLAP exclusively in liver-innervating cholinergic neurons. PLAP staining revealed the extensive cholinergic nerve fibers in the periportal and midlobular areas, further supporting the presence of hepatic cholinergic innervation.

Compared to the experimental groups, ChAT^Cre^ mice fed HFD showed elevated basal and fasting glucose levels and higher basal insulin levels, which led to hyperglycemic and hyperinsulinemic conditions during HFD feeding. These conditions caused hepatic steatosis in control mice. Interestingly, the deletion of parasympathetic cholinergic input to the liver reversed the detrimental effects of high-fat feeding on insulin sensitivity and hepatic fat accumulation.

The effects observed were partly due to the decrease in hepatic *de novo* lipogenesis and increase in fatty acid oxidation in the livers of ChAT^Cre^ mice, which lack DMV cholinergic innervation. In other words, when DMV cholinergic innervation to the liver was deleted, genes associated with lipogenesis were expressed at lower levels, while genes involved in fatty acid oxidation were expressed at higher levels, leading to reduced levels of triglycerides. Hence, our results suggest a significant role for parasympathetic cholinergic innervation in the regulation of hepatic lipid metabolism under obesogenic conditions.

Simple hepatic steatosis resulted from an imbalance between the delivery and production of fat in the liver and its subsequent secretion and metabolism [69]. Excessive dietary fat led to increased plasma concentrations of TGs and free fatty acids (FFAs), resulting in increased uptake into hepatocytes [70]. In our preparations, both groups consumed the same amount of food, implying that there may not be a substantial difference in the daily intake of fat. Additional sources of fatty acids can also be obtained from the breakdown of TGs stored in adipocytes, which is mainly controlled by insulin [71]. Insulin regulates systemic lipid metabolism by suppressing lipolysis in adipocytes. This mechanism is the primary way that insulin controls the breakdown of fats in these cells. As a result, insulin typically increases the uptake of FFAs from lipoproteins and decreases their release from adipocytes [70]. In contrast, insulin resistance depressed both of these functions [70]. ChAT^Cre^ mice, which lack DMV cholinergic neurons innervating the liver, displayed enhanced insulin sensitivity compared with the control group. This metabolic profile, including decreased adiposity, lower levels of plasma glycerol, and reduced fat mass, may contribute to the improved insulin sensitivity observed in experimental mice. By improving insulin resistance in adipocytes, the flow of FFAs and glycerol to the liver was limited, preventing the development of hepatic steatosis in ChAT^Cre^ mice that lack cholinergic innervation to the liver during HFD feeding.

We should highlight that disrupting the brain-liver axis caused the browning of ingWAT, which was a process that enhances glucose uptake and significantly contributes to systemic glucose disposal. This improvement in glucose disposal can ultimately lead to better insulin sensitivity [36]. Hepatocytes secrete various signaling molecules, such as bile acids, which can trigger WAT browning through TGR5 [12, 40, 72]. Daily administration of the TGR5 agonist for 1 week resulted in a significant decrease in subcutaneous white adipose tissue (scWAT) mass and the upregulation of browning marker genes, including *Ucp1* and *Cidea* [12]. ChAT^Cre^ mice lacking liver-projecting cholinergic neurons showed a significant reduction in WAT mass and adipocyte size. Interestingly, these mice also exhibited an increase in thermogenic gene and protein expression in ingWAT, leading to an elevation in nocturnal ingWAT temperature. The upregulation of *Cyp7b1* and bile acid levels due to the loss of hepatic cholinergic innervation may lead to browning of ingWAT as a result of crosstalk between the liver and ingWAT.

Communication between ingWAT and other organs, including the liver and muscle, is essential for the proper functioning of systemic metabolism [71]. Our current study revealed that the crosstalk between the liver and ingWAT through bile acids played a significant role in the progression of hepatic steatosis. Remarkably, silencing *Tgr5* in the ingWAT completely reversed the beneficial effects of the loss of hepatic cholinergic innervation. *Tgr5* knockdown resulted in enlarged adipocytes, leading to hypertrophy. Surprisingly, knockdown of *Tgr5* that was restricted to ingWAT induced lipid accumulation in the livers of ChAT^Cre^ mice lacking cholinergic innervation. In other words, the absence of hepatic cholinergic innervation failed to prevent the onset of hepatic steatosis following the silencing of *Tgr5* in ingWAT, suggesting that the ingWAT makes a crucial contribution to the development of hepatic steatosis. In fact, knockdown of *Tgr5* in ingWAT elevated plasma glycerol levels, indicating a role of TGR5 in regulating lipolysis in ingWAT. While our study provides evidence for crosstalk between the liver and ingWAT via TGR5, it is possible that TGR expressed in central neurons may also play a role in the observed effects as activation of hypothalamic TGR5 suppressed food intake and prevented diet-induced obesity [73, 74].

In conclusion, our study provides substantial evidence for the presence of parasympathetic cholinergic input to the liver, which constitutes a key component of the brain-liver axis. This axis is involved in the synthesis and secretion of bile acids, and the interorgan communication between the liver and ingWAT via bile acids plays a crucial role in the development of systemic insulin resistance and hepatic steatosis under obesogenic conditions. Therefore, targeting parasympathetic cholinergic input to the liver may offer a promising avenue for treating systemic insulin resistance and its associated hepatic steatosis in obesity.

## Materials and methods Ethics statement

All mouse care and experimental procedures were approved by the Institutional Animal Care Research Advisory Committee of the Albert Einstein College of Medicine (Approval number: 00001377) and performed in accordance with the guidelines described in the NIH Guide for the Care and Use of Laboratory Animals. Viral injections were performed under isoflurane anesthesia.

### Animals

Eight-nine-week-old ChAT^Cre^ mice were purchased from Jackson Laboratory (JAX stock # 006410)[75]. We backcrossed them to C57BL/6J mice for several generations, and the mutant mice were bred together to make this line homozygous. Mice were housed in cages under conditions of controlled temperature (22 °C) with a 12:12 h light-dark cycle and fed a high-fat diet (HFD; Research Diets, D12492; 20% calories by carbohydrate, 20% by protein, and 60% by fat) for 10-12 weeks, with water provided *ad libitum*. Mice were euthanized with an overdose of isoflurane at the end of the experiment.

### Western Blotting

Proteins were isolated from the liver using CelLytic^TM^ MT (Sigma, C3228) in the presence of a protease inhibitor/phosphatase inhibitor cocktail (Thermo Fisher Scientific, 78443). Proteins from the ingWAT were extracted using a protein extraction kit for adipocyte tissue (Invent, AT-022).

Protein concentrations were determined using the BCA Protein Assay Kit (Thermo Fisher Scientific, 23225). Proteins (30-50 ug) were prepared by adding laemmli sample buffer (Bio-Rad, 1610747) and heated at 95°C for 5min. The samples were then separated using 10% SDS-PAGE and transferred to a polyvinylidene fluoride (PVDF) membrane. The PVDF membrane was incubated with 5% w/v non-fat dry milk or 5% BSA for 1 h at room temperature and immunoblotted with anti-HSL (phospho-serine 660) (1:1000, Cell Signaling Technology, 45804S), anti-HSL (1:1000, Cell Signaling Technology, 4107), rabbit anti-UCP1(1:1000, Abcam, ab234430), rabbit anti-CHRM1 (1:200, Alomone Labs, AMR-001), rabbit anti-CHRM2 (Alomone Labs, AMR-002), rabbit anti-CHRM3 (Alomone labs, AMR-006), rabbit anti-CHRM4 (Alomone labs, AMR-004), rabbit anti-CHRM5 (Alomone Labs, AMR-005), and rabbit anti-β-actin (Cell Signaling, 4970S) antibodies. The specificity of anti-CHRM1-5 antibodies was tested by preincubation with the blocking peptide (CHRM1 with BLP-MR001, CHRM2 with BLP-MR002, CHRM3 with BLP-MR006, CHRM4 with BLP-MR004, and CHRM5 with BLP-MR005). Following incubation with primary antibodies, the membrane was washed three times in TBS-T and then incubated with an anti-rabbit IgG, HRP-linked antibody (1:10,000, Cell Signaling Technology, 7074) for 2 h at room temperature. ECL reagents were applied to the membrane and protein bands were detected using an Odyssey Fc imaging system (Li-COR) and ChemiSOLO (Azure biosystems).

### Immunostaining to examine the hepatic cholinergic system

To examine DMV cholinergic neurons innervating the liver, ChAT^Cre^ mice received an AAV.Php.eB-flex-tdTomato (titer; 1.9x10^13^ pfu/ml, Addgene 28306-PHP.eB) injection into the liver. We also injected AAV-phSyn1(s)-FLEX-tdTomato-T2A-SypGFP-WPRE (Addgene 51509- AAV1) into the DMV (500 nl of 4 x 10^12^ pfu/ml per site; coordinate; AP: -7.5 mm, ML: ±0.1 mm, DV: -3.5 mm) to label cholinergic nerve terminals in the liver. Mice were killed 5 and 10 weeks post-viral inoculation.

Mice were anesthetized with isoflurane (3%) and transcardially perfused with pre-perfusion solution (9 g NaCl, 5 g sodium nitrate, 10,000 U heparin in 1L distilled water) followed by 4% paraformaldehyde solution. Liver sections were stained with a rabbit anti-GFP antibody (Rockland, 600-401-215, 1:500) to visualize cholinergic nerve terminals. Brainstem sections were labeled with rabbit anti-RFP (Rockland, 600-401-379, 1:500) and goat anti-ChAT antibodies (Millipore, AB144P, 1:200). The expression of muscarinic ACh receptors CHRM2 and 4 was examined using rabbit anti-CHRM2 (Alomone, AMR-002, 1:200) and rabbit anti-CHRM4 (Alomone, AMR-004, 1:200) antibodies. Cholinergic nerve terminals were stained with goat anti-vAChT (Millipore, ABN100) and goat anti-ChAT (Millipore, AB144P) antibodies. The tissue samples were incubated with AlexaFluor 568 donkey anti-goat IgG (Invitrogen, A11057, 1:200), AlexaFluor 568 donkey anti-rabbit IgG (Invitrogen, A11011, 1:200), and AlexaFluor 488 donkey anti-rabbit IgG (Invitrogen, A21206, 1:200). The tissues were washed, dried, and mounted with VECTASHIELD medium containing DAPI. Images were acquired using a Leica SP8 confocal microscope.

To visualize cholinergic nerve fibers in the liver, ChAT^Cre^ mice were given an injection of AAVrg-CMV-FLEX-PLAP (with a titer of 1.0x10^12^ GC/ml from ABM, Inc.) into the liver. After two months following viral infections, the mice were perfused with PBS and fixed with a 4% paraformaldehyde solution. The liver tissues were then sliced to a thickness of 30 μm using a vibratome and incubated in alkaline phosphatase (AP) buffer (0.1 M Tris HCl pH 9.5, 0.1 M NaCl, 50 mM MgCl2, 0.1% Tween20, and 5 mM levamisole) for one hour and thirty minutes at room temperature. The tissue was then visualized using the NBT/BCIP substrate solution (ThermoFisher Scientific, 34042) until the desired stain developed. The tissues were washed with distilled water and mounted with aqueous mounting media (Vector, H-5501). The stained samples were captured in brightfield scanning mode using the Panoramic 250 FLASH III F2, and analyzed using slideViewer 2.7 (3DHISTECH).

### Clear, unobstructed brain imaging cocktails and computational analysis (CUBIC) tissue clearing of the brainstem and liver tissues

The CUBIC method [76] was used to clear liver and brainstem tissues. Mice were perfused with 4% (w/v) paraformaldehyde (PFA) in PBS. Liver and brain tissues were collected and post-fixed in 4% PFA at 4C° for 24 h. Tissue samples were sectioned using a vibratome at a thickness of 300 μm. The sections were immersed in CUBIC reagent 1(25wt% Urea, 25wt% Quadrol, 15wt% Triton X-100) with gentle shaking at 37C° for 3 days. After decolorization (transparent tissues), the sections were washed with PBS three times at RT for 2 h and then incubated in a blocking solution composed of 1% donkey serum, 0.5% BSA, 0.3% Triton X-100, and 0.05% sodium azide for 3 h at RT with rotation. The tissue samples were incubated with goat anti-ChAT (Millipore, AB144P, 1:200), rabbit anti-RFP (Rockland, 200-301-379 1:200), and rabbit anti-GFP (Rockland, 600-401-215, 1:200) antibodies for 5 days at RT with mild shaking, and washed three times with PBS (each for at least 2 h). Following wash-out, the tissue samples were incubated with AlexaFluor 568 donkey anti-goat IgG (1:200, Invitrogen, A11057), AlexaFluor 568 donkey anti-rabbit IgG (1:200, Invitrogen, A11011), and AlexaFluor 488 donkey anti-rabbit IgG (1:200, Invitrogen, A21206) for 3 days at room temperature. Tissues were washed and immersed in CUBIC-reagent 2 (50wt% Sucrose, 25wt% Urea, 10wt% Triethanolamine, and 0.1%(v/v) Triton X-100) for 1 d and mounted with a mixture of mineral oil and silicone oil. A z-stack consisting of 120 images was acquired with 0.33 μm steps with a Leica SP8 confocal microscope, resulting in a total optical thickness of approximately 40 μm and 3D construction was performed with AIVIA (version 13.1).

### Viral injection to delete DMV cholinergic neurons innervating the liver

pAAV-mCherry-FLEX-dtA (Addgene, 58536) was subcloned and packaged in an AAV serotype retrograde (Applied Biological Materials, Inc.). A total volume of 20μl of AAVrg-mCherry-FLEX-dtA (titer, >1 × 10^12^ pfu/ml) was injected into the medial and left lobes of the livers of the ChAT^Cre^ mice. AAVrg-FLEX-tdTomato viruses (titer, >1x10^13^ pfu/ml) were used as the controls.

To quantify cell death over time, the mice were killed 10-12 weeks after viral inoculation. Mice were anesthetized with isoflurane (3%) and transcardially perfused with PBS and heparin, followed by 10% neutral buffered formalin solution (Sigma, HT501128). Brainstem and liver tissues were embedded in paraffin to create paraffin blocks. Paraffin blocks were cut into 10 µm sections for the brainstem and 5 µm sections for liver tissues at our histology and comparative pathology core facility.

Coronal sections of the brainstem from approximately −7.2 to −7.6 mm posterior to bregma were stained with an anti-ChAT antibody (Millipore, AB144P). Cell counting of DMV cholinergic neurons was performed using the ImageJ software (version FIJI). The adipocyte cell area was measured using the Adiposoft plugin (version 1.16) in the Image J software.

We administered *Tgr5* siRNA or control siRNA (Titer, 1X10^9^ IU/ml, volume 20 μm) into the ingWAT of ChAT^Cre^ mice without liver-innervating cholinergic neurons to exclusively knock down *Tgr5* in the ingWAT.

### Measurement of body weight, body composition, and blood glucose

Body weight was measured weekly at 9 am. Body composition for fat mass and fat-free mass was assessed by ECHO MRI at our animal physiology core. Blood samples were collected from the mouse tail, and a small drop of blood was placed on the test strip of a glucose meter. Non-fasting basal glucose levels were measured at 9:00 am. Fasted blood glucose levels were measured after overnight fasting at 10 weeks post-viral injection.

### Assessment of energy expenditure and locomotor activity

To examine if the loss of function of hepatic cholinergic input regulates energy expenditure and locomotor activity, we performed indirect calorimetry on mice fed HFD for 10 weeks. Mice were individually housed in the calorimeter cages and acclimated to the respiratory chambers for at least 2 days prior to gas exchange measurements. Indirect calorimetry was performed for 3 days at the end of 10 weeks using an open-circuit calorimetry system. O_2_ consumption and CO_2_ production were measured for each mouse at 10-min intervals over a 24- h period. The respiratory exchange ratio was calculated as the ratio of CO_2_ production over O_2_ consumption. Since there was no difference in food intake, VO_2_ was normalized for body lean mass. Locomotor activity in X-Y and Z planes was measured by infrared beam breaks in the calorimetry cages.

### Assessment of glucose tolerance and insulin tolerance

For GTT, the experimental and control mice were fasted for 15 h (6:00 pm–9:00 am) at 10 weeks post-viral inoculation. Sterile glucose solution was intraperitoneally administered at a concentration of 2 g/kg (glucose/body weight) at time 0. Blood glucose levels were measured at 15, 30, 60, 90, and 120 min after glucose injection. Blood glucose levels versus time after glucose injection were plotted, and the area under the curve was calculated and compared between the experimental and control groups.

For ITT, the mice were fasted for 4 h (9:00 am to 1:00 pm). Blood glucose levels were measured at 0, 15, 30, 60, 90, and 120 min following i.p. injection of insulin (1 U/kg). We immediately injected glucose (2 g/kg) if the mice appeared ill owing to insulin-induced hypoglycemia.

### RT-qPCR analysis

Liver tissues were collected from the experimental and control groups 10-12 weeks post viral inoculation. Liver tissues were homogenized in Trizol reagent (ThermoFisher Scientific, 15596-018), and total RNAs were isolated according to the manufacturer’s instructions. First-strand cDNAs were synthesized using the SuperScript III First-Strand synthesis kit (ThermoFisher Scientific, 18080-051). qPCR was performed in sealed 96-well plates with SYBR Green I master Mix (Applied Biosystems, A25742) using the Quant Studio 3 system (Applied Biosystems). qPCR reactions were prepared in a final volume of 20 μl containing 2 μl cDNAs, and 10 μl of SYBR Green master mix in the presence of primers at 0.5 μM. Glyceraldehyde 3- phosphate dehydrogenase *(Gapdh)* for liver and 18S ribosomal RNA (*18s rRNA*) for white adipocyte tissue were used as an internal control for quantification of each sample. Amplification was performed under the following conditions: denaturation at 95 °C for 30 seconds, followed by 40 cycles of denaturation at 95 °C for 30 seconds, and annealing/extension at 60 °C for 1 minute.

We examined the following genes; *Srebp1c* (NM_001358315.1), *Chrebp* (NM_021455.5), *Acly* (NM_134037.3)*, Acaca* (NM_133360.3)*, Fasn* (NM_007988.3)*, Scd1* (NM_009127.4)*, Ppara* (NM_011144.6)*, Prdm16* (NM_027504), *Cpt1a* (NM_013495.2)*, Cpt2* (NM_009949.2)*, Acadm* (NM_007382.5)*, Acadl* (NM_007381.4)*, G6pase* (NM_008061.4)*, Pck1* (NM_011044.3)*, Slc2a2* ( NM_031197.2)*, Pgc1a* ( NM_008904.3)*, Ucp1* (NM_009463.3)*, Dio2* (NM_010050.4)*, Cidea* (NM_007702.2)*, Gpbar1* (NM_174985.1)*, Cyp7a1* (NM_007824)*, Cyp7b1* (NM_007825) (S1 Table). All the primer sequences will be validated with “Primer BLAST” primer design program to ensure specificity for the target gene. Melt curves and gel electrophoresis were analyzed to confirm the specificity of the PCR products. Relative gene expression was determined using a comparative method (2^-ΔΔCT^).

**Table 1.**
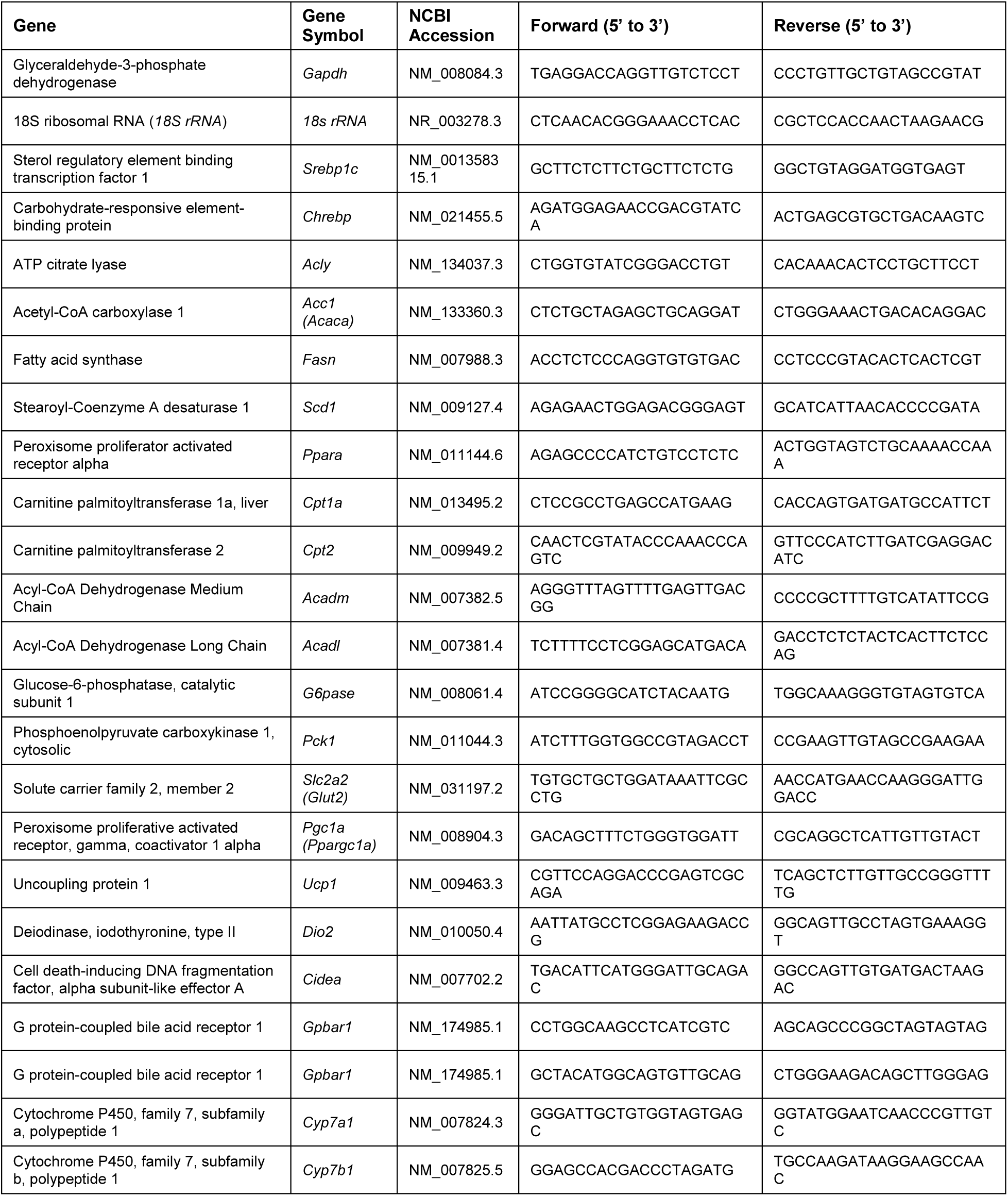
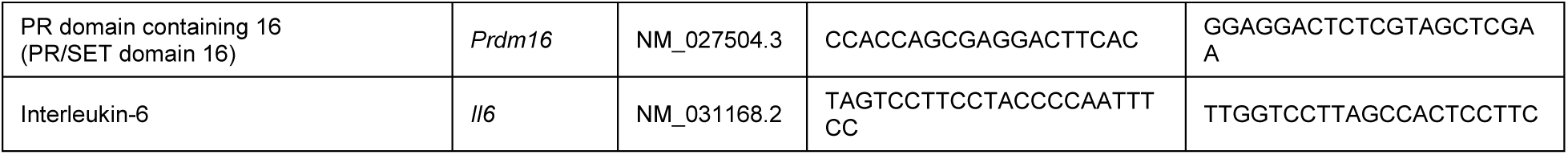
List of primer sets for qPCR

### Spatial Transcriptomics

Livers were snap-frozen in liquid nitrogen and thawed briefly on ice, followed by embedding in a cryomold (Sakura, 4565) filled with Neg-50 (Epredia, 6502). The cryomold was then frozen by placing into a container with pre-chilled 2-methylbutane on dry-ice. The samples were cryosectioned (Cryostar NX70 Cryostat, -18C for blade, -18 °C for specimen head, set at 10mm thickness) and placed on a pre-chilled Visium Spatial Gene Expression Slide (10x Genomics, PN-2000233). The tissue sections were processed following 10x Genomics protocols (PN-1000187, PN-1000215). The adaptors and sequencing primers were obtained from Illumina. The libraries were sequenced by Novogene using the NovaSeq 6000 (PE150) instrument targeting 80,000 reads per spot. Fastq files were aligned against mouse mm10 reference and histology images were processed with Spaceranger (2.0). Pericentral, mid-zone and periportal clusters were identified based on Graph-Based clusters and known zonation markers (*Cyp2e1*, *Glul*, *Gulo* for pericentral, *Cyp2f2*, *Hsd17b13*, *Sds* for periportal). Violin plots were generated using Loupe Browser (6.4.1).

### Measurement of hepatic and plasma TG and ACh levels

Liver samples were harvested from mice with or without liver-innervating cholinergic neurons. Tissue samples were weighed and homogenized in 1 mL of diluted NP-40 solution to solubilize triglycerides. The samples were centrifuged at 16,000 × g to remove insoluble material. The TG levels were measured using a TG assay kit (Cayman Chemical, 10010303).

For the acetylcholine assay, liver tissues were homogenized in PBS. The homogenates were then centrifuged at 5,000 g for 5 min. The supernatants were collected, and ACh levels were determined using an acetylcholine ELISA kit (Novus, NBP2-66389).

### Measurement of plasma insulin levels and fasting serum bile acid levels

Blood samples from the experimental and control groups were collected from the retroorbital plexus using heparinized capillary tubes. Whole blood was centrifuged at 12,000 × g for 10 min, and plasma was separated and stored at -20°C until use. Plasma insulin concentrations were determined using two-site sandwich ELISA kits (Mercodia, cat# 10-1247- 0). To analyze fasting total bile acid levels in the serum, fasting blood samples were collected without anticoagulant and determined using mouse total bile acid (Crystal Biochem, Cat#80471). Bile acid profiling was performed at our metabolomic core facility.

### Statistics

All statistical results are presented as the mean ± SEM. Statistical analyses were performed using GraphPad Prism, version 10.0. Two-tailed *t*-tests were used to determine the p-values for each pairwise comparison. The Mann-Whitney U test was performed to evaluate the differences in bile acid composition between the groups. Time-course comparisons between groups were analyzed using a two-way repeated-measures analysis of variance (RM ANOVA) with Sidak’s correction for multiple comparisons. Data were considered significantly different when the probability value was less than 0.05.

## Data availability

The spatial transcriptomic dataset was submitted to GEO and assigned the accession number GSE255552.

## Acknowledgments

**General**: We thank Drs. Streamson Chua Jr. and Fajun Yang for their valuable feedback and comments on this study. We also thank Dr. Shun-Mei Liu and Licheng Wu for their technical assistance.

**Funding:** This work was supported by the NIH (R01 AT011653 and R03 TR003313 to Y.-H.J, R01 DK092246 to Y.-H.J and G.J.S., P30 DK020541 to Y.-H.J, G.J.S., and J.E.P. and DK110063 to J.E.P.).

**Author contributions:** J.Y.H performed viral injection, immunostaining, Western blotting, and ELISA assays, and analyzed data. J.O. and L.L. performed and analyzed spatial transcriptomics analysis. Y.-H.J designed the research, performed viral injection and immunocytochemistry, analyzed the data, and wrote the manuscript with inputs from G.J.S. and J.E.P. G.J.S and J.E.P. also designed the study.

**Competing interests:** The authors declare no conflict of interest.

## Supplementary Figure legends

**S1 Fig.** Parasympathetic cholinergic system in the mouse liver.

(A) Graph showing mRNA expression of *Chrm1, 2, 3, 4,* and *5* in mouse liver samples.

(B) Images of Western blot analysis of liver homogenates showing the expression of CHRM1-5, except for CHRM3.

(C) Western blot images showing the expression of CHRM3 in the brain, but not the liver samples. The antibodies used in this study were Alomone Labs (AMR 006) and Abcam (ab87199).

(D) Images of confocal fluorescence microscopy showing the expression of CHRM4 and vAChT in the liver parenchyma (upper panel). Scale bar, 25 μm, Bottom panel: Higher magnification view of the white dotted square. CHRM4s were found adjacent to vAChT-positive nerve terminals (white arrows). Blue: nucleus staining with DAPI. Scale bar, 10 μm

(E) Images of confocal fluorescence microscopy showing the expression of CHRM2 in hepatocytes. The vAChT-positive parasympathetic nerve terminals were adjacent to CHRM2. Scale bar: 50 μm Bottom panel: Higher magnification view of the white dotted square. Arrowheads represent examples of cholinergic synapses on hepatocytes The data supporting the graphs shown in the figure (Fig. S1A) are available in the S2 Data file.

**S2 Fig.** Ablation of DMV cholinergic neurons innervating the liver using AAVrg-FLEX-dtA.

(A and B) Graphs showing the number of cholinergic neurons on the left and right sides of the DMV in the control and experimental groups. Unpaired *t*-test, *p<0.05; ***p<0.001 The data supporting the graphs shown in the figure (Fig. S2A and B) are available in the S2 Data file.

**S3 Fig.** Expression patterns of liver zonation markers in controls fed a standard chow diet.

(A) Image of confocal fluorescence microscopy showing the expression of E-Cad and GS in the liver parenchyma. Scale bar, 100 μm

**S4 Fig.** Ablating parasympathetic cholinergic innervation to the liver in ChAT^Cre^ female mice does not cause hepatic lipid accumulation.

(A) Macroscopic appearance of the livers of ChAT^Cre^ female mice receiving AAVrg-FLEX-GFP and AAVrg-FLEX-dtA. E-Cad (green) and GS (red) staining in the control and the experimental groups. Scale bar, 100 μm

(B and C) H&E and Oil Red O staining of liver tissues from the control and experimental groups. No histological differences were observed between the groups (upper panels, scale bar, 50 μm; bottom panels, scale bar, 100 μm)

**S5 Fig.** Deleting parasympathetic cholinergic input to the liver does not alter the expression of gluconeogenic enzyme genes.

(A) Relative *Il6* mRNA expression in the livers of the control (n= 7 mice) and experimental (n = 7 mice) groups. Unpaired *t*-test, **p<0.01

(B) Graph showing mRNA expression of the gluconeogenic enzymes in the livers of the control and the experimental groups (control, n=8 mice; experimental, n=7 mice). The data supporting the graphs shown in the figure (Fig. S5A and B) are available in the 2S2 Data file.

**S6 Fig.** No changes in sympathetic innervation of the liver in ChAT^Cre^ mice without parasympathetic cholinergic innervation.

(A and B) Images of confocal fluorescence microscopy showing TH-positive nerve fibers in the liver parenchyma of ChAT^Cre^ mice with and without liver-projecting cholinergic neurons (arrows). Scale bar, 30 μm. Right panel: Higher magnification view of the white dotted square

(C) Graph showing hepatic norepinephrine levels in the control (n=8 mice) and experimental (n=8 mice) groups. The data supporting the graphs shown in the figure (Fig. S6C) are available in the S2 Data file.

S7 Fig. Differential gene expression in the livers of mice with and without liver-innervating parasympathetic cholinergic neurons.

(A) Violin plots depicting the differential expression of the liver zonation marker genes in the pericentral, midlobular, and periportal areas in the control and experimental groups.

(B) Volcano plots illustrate the differences in gene expression across the zonation of the livers in control and experimental mice.

(C) Violin plots displaying the enriched genes across the zonation of the livers in control and experimental mice.

(D) Plots displaying the top GO terms in biological process and mammalian phenotype across the zonation of the livers in control and experimental mice. The data supporting the graphs shown in the figure (Fig. S7A-D) are available in the S2 Data file.

**S8 Fig**. Ablating parasympathetic cholinergic innervation to the liver in ChAT^Cre^ mice does not impact food intake but does increase energy expenditure.

(A) Graph showing changes in body weight of the control and experimental groups during high-fat diet feeding. There was a significant difference in body weight gain between the groups (control, n= 12 mice; experimental mice, n=21 mice; two-way ANOVA test, *p<0.05).

(B) Graphs show a significant difference in body fat but not fat-free mass between the two groups (fat mass and lean mass, control, n= 9 mice; experimental mice, n=10 mice, unpaired *t*- test, *p<0.05).

(C) Graphs showing that the experimental mice (n=7) exhibited a significantly lower ingWAT mass than the control group (n=8), whereas there was no difference in liver, interscapular WAT (isWAT), gonadal WAT (gWAT) and perigonadal WAT (pgWAT) mass between the groups. Unpaired *t*-test, *p<0.05

(D) Graphs showing that both groups consumed the same amount of food (control, n= 5 mice; experimental mice, n= 7 mice).

(E and F) Summary plots showing VO_2_ and VCO_2_ between the groups. A significant difference in VO_2_ and VCO_2_ in the dark phase was observed between the groups (control, n = 7 mice, experimental mice, n= 8 mice). Unpaired *t*-test, *p<0.05

(G and H) Summary plots showing total energy expenditure (TEE) and respiratory exchange ratio (RER) between the groups (control, n = 7 mice, experimental mice, n= 8 mice). The loss of hepatic cholinergic input significantly increased TEE in the dark phase. Unpaired *t*-test, *p<0.05 (I and J) Summary plots showing total locomotor and ambulatory activities between the experimental groups (control, n = 7 mice, experimental mice, n= 8 mice). Unpaired *t*-test, *p<0.05 The data supporting the graphs shown in the figure (Fig. S8A-J) are available in the S2 Data file.

**S9 Fig.** Enhanced insulin sensitivity in ChAT^Cre^ mice without parasympathetic cholinergic innervation to the liver.

(A and B) Summary graphs showing the basal (non-fasting) and fasting glucose levels in the control (open circle) and experimental (closed circle) groups. Unpaired *t*-test, *p<0.05

(C and D) Summary graphs showing the i.p. GTT in ChAT^Cre^ mice with (n= 5 mice) and without (n= 6 mice) liver-projecting cholinergic neurons. AUC: area under the curve

(E) Plot showing the i.p. ITT in ChAT^Cre^ mice with and without parasympathetic cholinergic neurons innervating the liver (Two-way ANOVA followed by Sidak multiple comparisons test, control, n=6 mice; experimental group, n=6 mice, F(1, 10) = 6.6, *p=0.03).

(F) Graph showing plasma insulin levels in the controls (n= 8 mice) and the experimental group (n= 7 mice). There was a significant difference in plasma insulin levels between the groups. Unpaired *t*-test, *p<0.05 The data supporting the graphs shown in the figure (Fig. S9A-F) are available in the S2 Data file.

**S10 Fig.** Ablating parasympathetic cholinergic innervation to the liver in ChAT^Cre^ female mice does not alter food intake or body weight.

(A) Graph showing changes in body weight of female ChAT^Cre^ mice with and without parasympathetic cholinergic innervation to the liver during high-fat feeding. There was no significant difference in body weight gain between the groups (control (open circle), n= 10 mice; experimental (closed circle), n=11 mice).

(B and C) Plots show no differences in body fat and lean mass between the two groups (control, n= 10 mice; experimental mice, n=11 mice).

(D) Graphs showing that both groups consumed the same amount of food (control, n= 6 mice; experimental mice, n= 7 mice).

(E and F) Summary graphs showing the basal (non-fasting) and fasting glucose levels. There were no significant differences in basal and fasting glucose levels between groups (control, n= 8 mice; experimental mice, n=9 mice)

(G and H) Summary graphs showing the i.p. GTT and i.p. ITT in mice with and without liver-projecting cholinergic neurons (GTT: control, n= 8 mice; experimental mice, n=9 mice; ITT: control, n= 9 mice; experimental mice, n=10 mice). No significant differences were observed between the groups. The data supporting the graphs shown in the figure (Fig. S10A-H) are available in the S2 Data file.

**S1 movie.** 3D projection view of confocal fluorescence microscopy images of hepatocytes receiving tdTomato-positive nerve terminals in cleared liver tissue of ChAT^Cre^;ChR2-tdTomato mice.

**S2 Movie.** 3D projection view of confocal fluorescence microscopy images showing the expression of GFP-positive nerve terminals in cleared liver tissue collected from ChAT^Cre^ mice injected with AAV1-FLEX-tdTomato-T2A-sypGFP as shown in Fig. 1H.

## Notes

### Competing Interest Statement

The authors have declared no competing interest.

### Summary of Updates

This revised version of the manuscript has been revised to add new results.

