## Supplementary figures and images for "The development of hepatic steatosis depends on the presence of liver-innervating parasympathetic cholinergic neurons in mice fed a high-fat diet"

### Supplementary figures

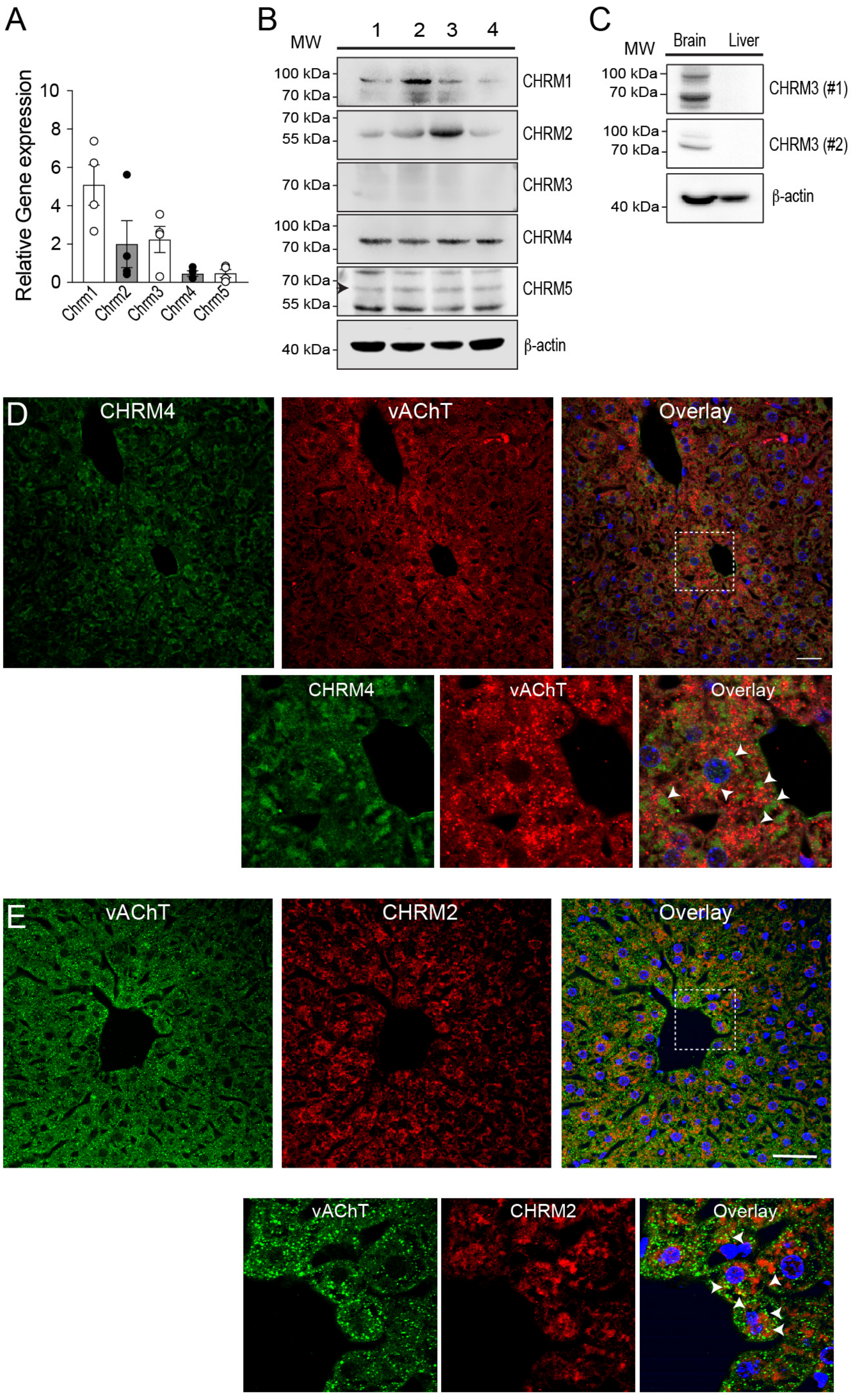

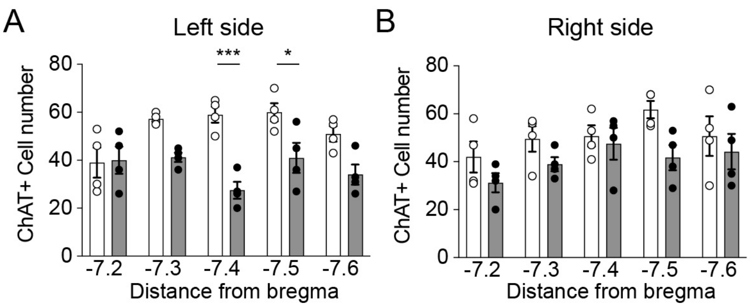

A. ChAT[Cre] fed a standard chow diet

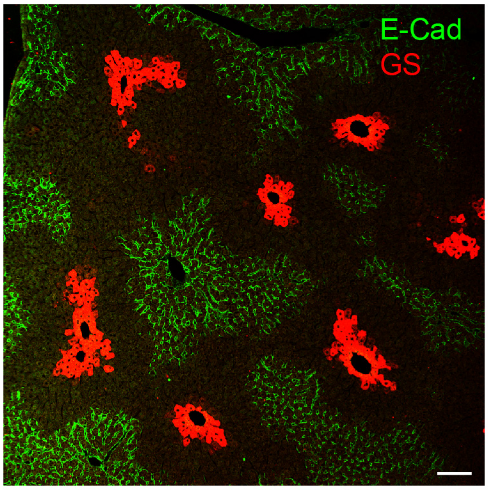


S3. Figure

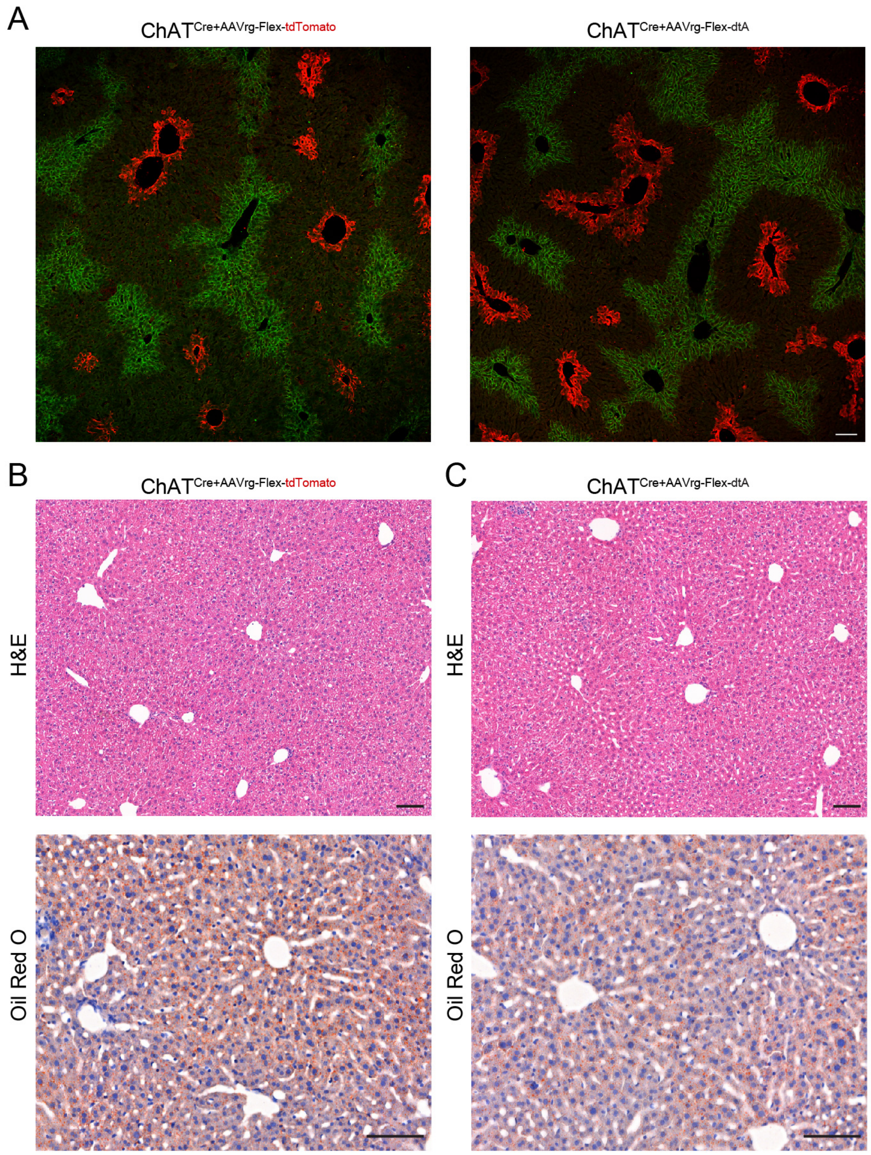

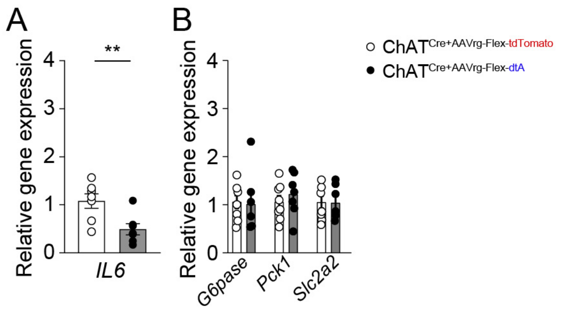

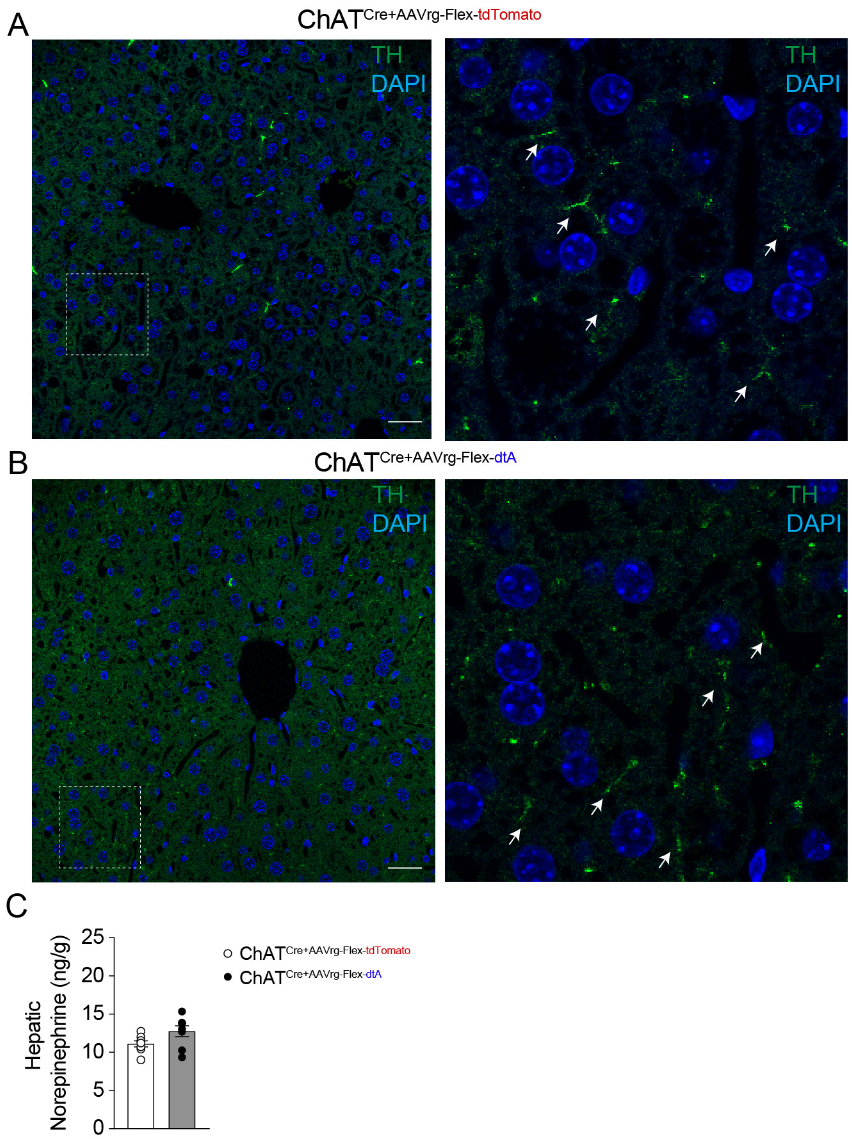

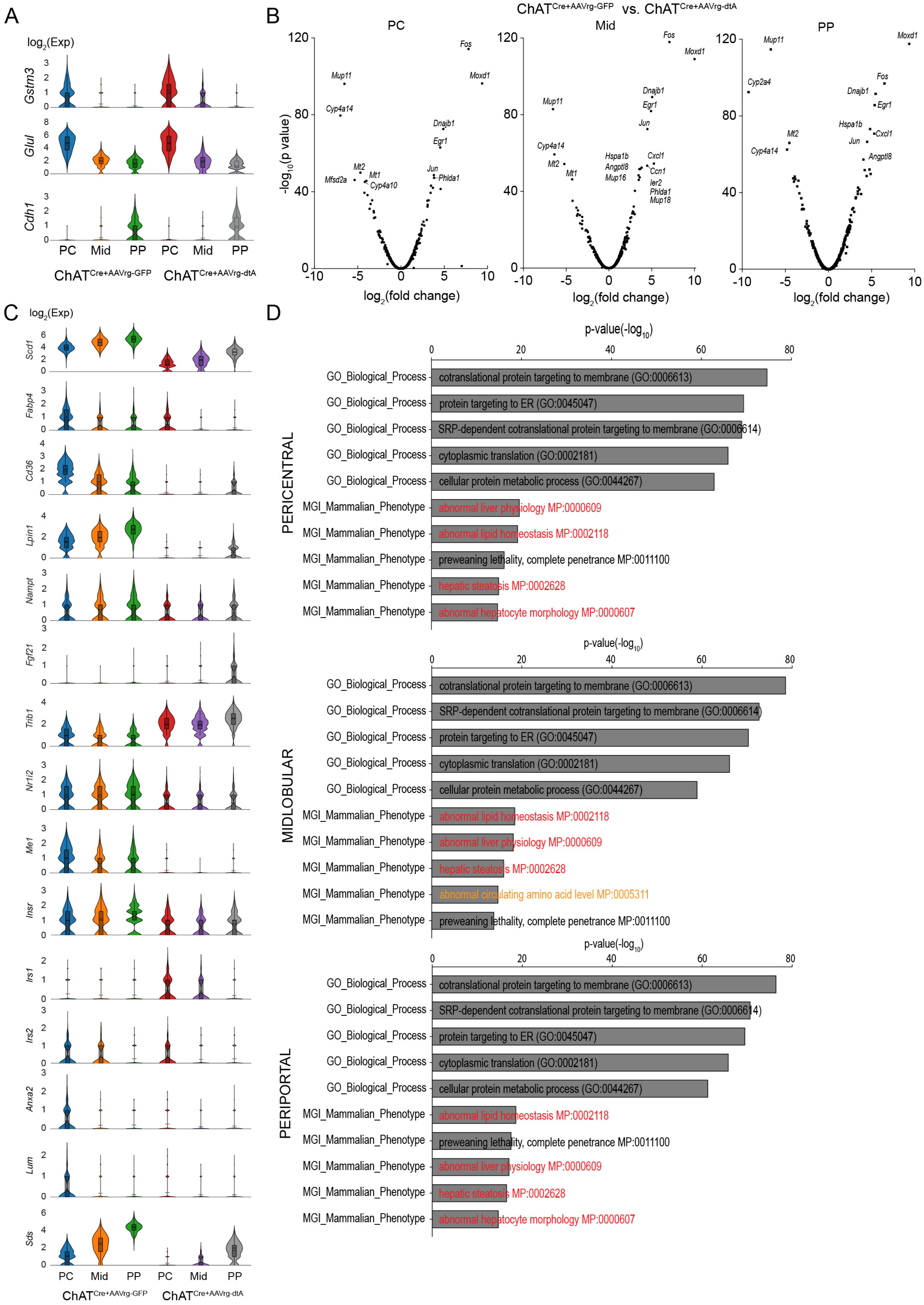

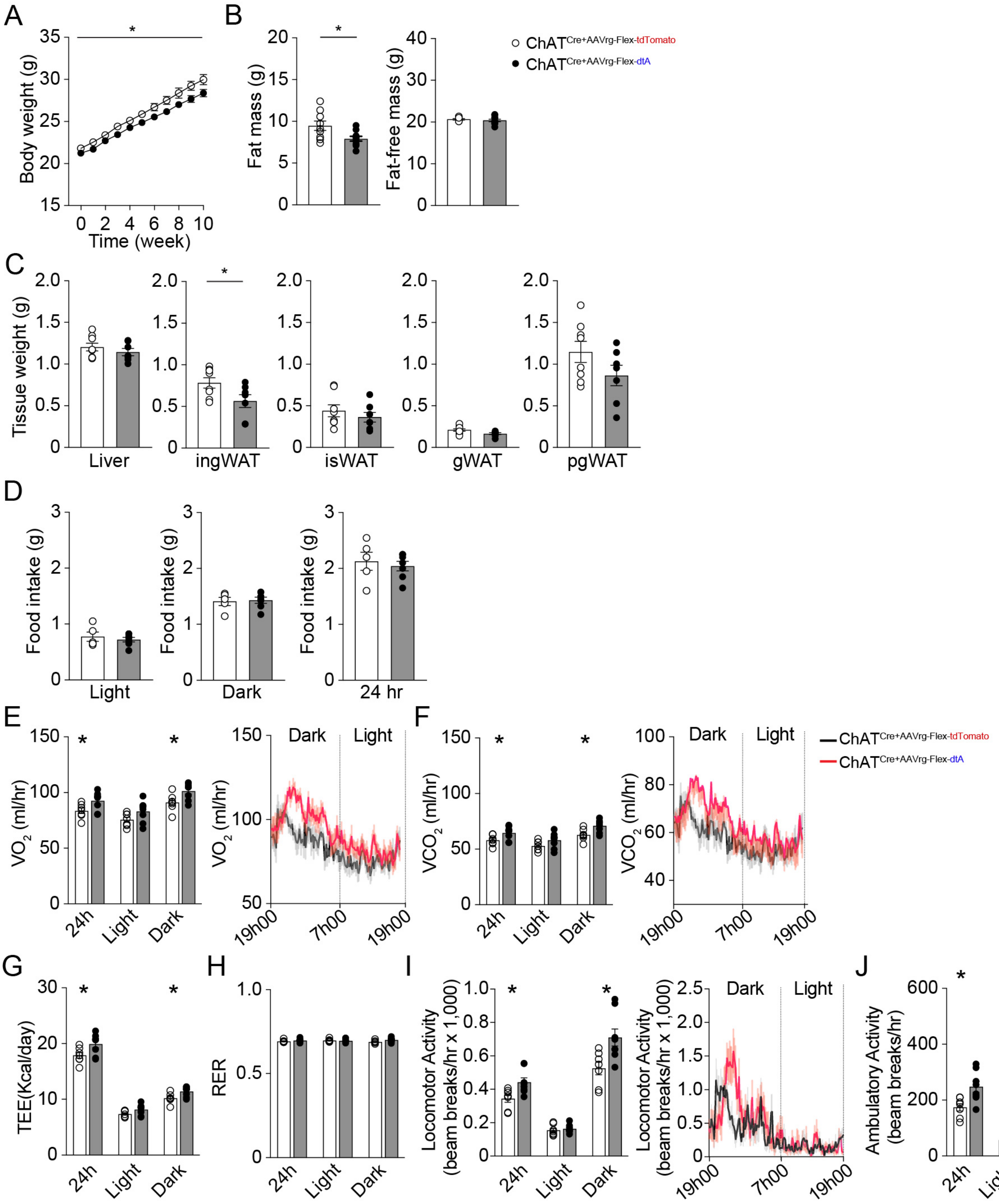

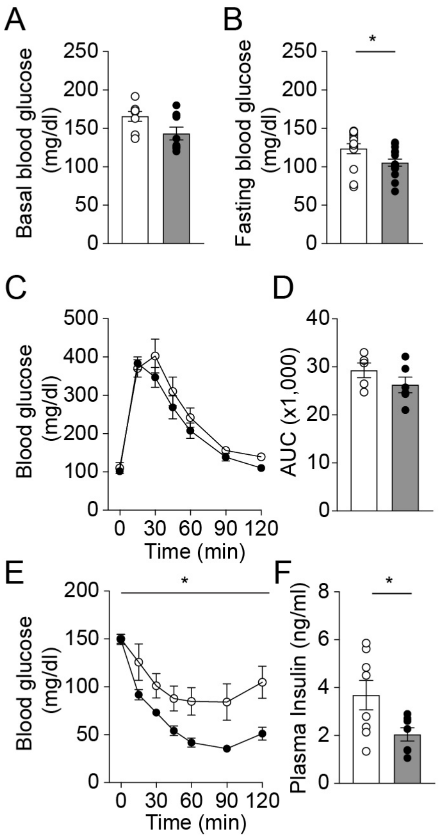

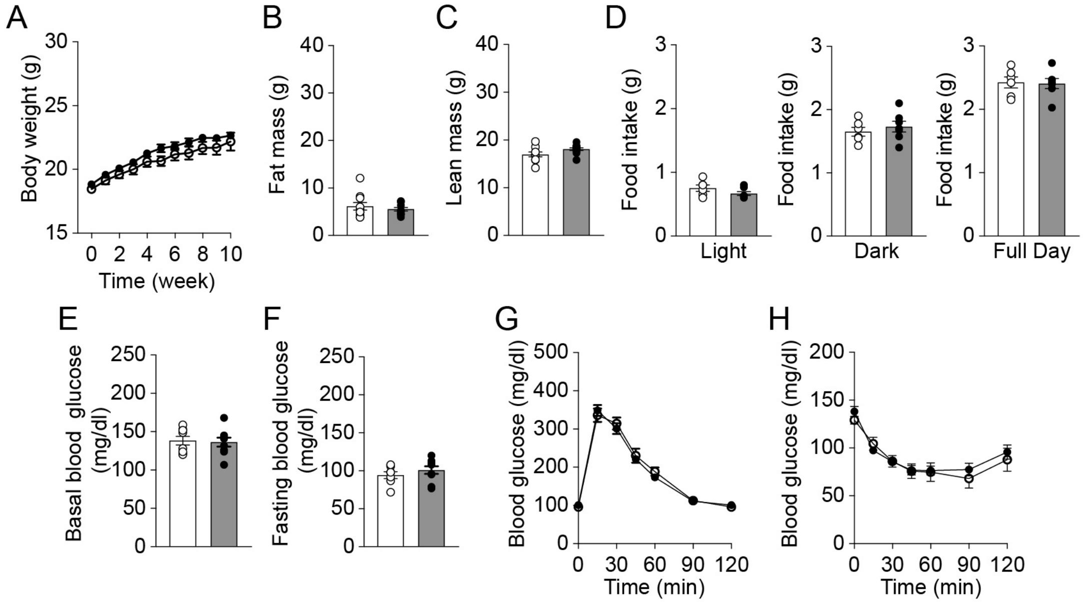
